# Selective innervation of subpopulations of striatal neurons by distinct sets of neurons of the external globus pallidus

**DOI:** 10.1101/2025.10.06.680630

**Authors:** Natalie M Doig, Calum McIntyre, Kouichi Nakamura, Andrew Sharott, Konstantinos Meletis, Peter J Magill

## Abstract

The striatum, the primary input nucleus of the basal ganglia, contains diverse populations of projection neurons and interneurons, each with distinct roles in motor processing. The external globus pallidus contains distinct populations of inhibitory neurons that are highly interconnected with other regions of the basal ganglia. How pallidal neurons innervate different populations of striatal neurons is critical for our understanding of their role in motor processing. Here we use monosynaptic viral tracing in transgenic mice to quantitively define the organization of pallidostriatal connectivity. We show that FoxP2-expressing arkypallidal neurons provide the greatest input to the striatum, primarily targeting indirect pathway spiny projection neurons and cholinergic interneurons. In contrast, prototypic pallidal neurons, defined by Nkx2-1 expression, preferentially innervate parvalbumin- and somatostatin-expressing interneurons. These data reveal a structured and selective organization of pallidostriatal projections, suggesting that distinct classes of pallidal neurons modulate striatal circuits through complementary pathways. Mapping these connections is key to understanding how basal ganglia networks coordinate motor control.

## Introduction

The external globus pallidus (GPe) is highly interconnected within the basal ganglia, a collection of subcortical nuclei critical for normal motor function. The GPe is uniquely positioned in that it innervates all other nuclei of the basal ganglia and is thus poised to have a significant effect on all aspects of basal ganglia function. Recent work has shifted from the view that the GPe is part of a linear pathway, cortex-striatum-GPe-basal ganglia output nuclei, towards the understanding that the GPe influences processing via reciprocal connections with all other basal ganglia nuclei (Courtney, Pamukcu and Chan 2023). The pallidostriatal pathway has been identified in a variety of species (Staines, Atmadja and Fibiger 1981; Beckstead 1983; Bevan *et al*. 1998; Kita, Tokuno and Nambu 1999; Bolam *et al*. 2000; Kita and Kita 2001), but the precise details of this pathway remain to be elucidated; in a large part due to the heterogeneity of both pallidal and striatal neurons (Hegeman *et al*. 2016).

At least two major cell types in the GPe have been characterised, referred to as prototypic and arkypallidal neurons (Mallet *et al*. 2012; Abdi *et al*. 2015; Dodson *et al*. 2015; Abecassis *et al*. 2020; Aristieta *et al*. 2021; Cui *et al*. 2021b). Prototypic neurons, which express homeobox protein Nkx2-1, and constitute the majority of GPe neurons (∼70-75%), innervate downstream (e.g., substantia nigra pars reticulata, SNpr) and upstream targets (striatum) (Abdi *et al*. 2015; Dodson *et al*. 2015). In contrast, arkypallidal neurons, which express the transcription factor FoxP2, only send projections to the striatum (Mallet *et al*. 2012; Dodson *et al*. 2015; Hegeman *et al*. 2016). Prototypic neurons can be further subdivided on the basis of the expression of other markers; most prototypic neurons also express parvalbumin (PV), these neurons are the classical GPe neurons with axonal arborizations in the subthalamic nucleus (STN), globus pallidus interna (GPi) and the SNpr (Abdi *et al*. 2015; Dodson *et al*. 2015). Prototypic and arkypallidal neurons can be further distinguished on the basis of firing characteristics, developmental origin, morphology, and activity during motor tasks (Mallet *et al*. 2012; Abdi *et al*. 2015; Dodson *et al*. 2015; Abecassis *et al*. 2020; Aristieta *et al*. 2021; Cui *et al*. 2021b), we hypothesize that the distinctive properties of GPe neurons are underpinned by distinctive innervation from striatal cell types including direct and indirect pathway spiny projection neurons (SPNs) and the three major populations of interneurons: GABAergic interneurons expressing either parvalbumin (PV) or somatostatin (SOM) and neuronal nitric oxide synthase (nNOS) and cholinergic interneurons expressing choline acetyltransferase (ChAT) (Sharott *et al*. 2012; Kondabolu *et al*. 2023).

The distinct populations of both striatal and GPe neurons have distinct roles in information processing and, movement control (Dodson *et al*. 2015; Mallet *et al*. 2016). However, in order to precisely understand the roles of the pallidostriatal network in behaviour it is critical to understand the pattern of innervation of subpopulations of striatal neurons by sub-populations of pallidal neurons. To address this we employed a combination of genetic and viral techniques; using a retrograde modified rabies virus in concert with a variety of transgenic mouse lines to selectively target distinct populations of the GPe and striatum neurons (Pollak Dorocic *et al*. 2014; Ährlund-Richter *et al*. 2019; Kondabolu *et al*. 2023). Using design-based stereology, we estimated connectivity between specific cell types of the pallidum and striatum. Arkypallidal neurons preferentially innervate striatal indirect pathway neurons and cholinergic interneurons, whereas prototypic pallidal neurons preferentially innervate direct pathway projections neurons and the GABAergic interneurons. Our findings reveal that pallidostriatal connectivity is not diffuse but highly selective, establishing a structural framework through which the GPe can exert targeted control over striatal output and, by extension, behaviour.

## Results

### Quantification of neuronal cell types constituting the pallidostriatal pathway

To identify and quantify molecularly characterized neurons of the GPe innervating neuronal populations of the dorsal striatum we employed a retrograde monosynaptic viral tracing strategy in concert with transgenic mouse lines expressing Cre-recombinase in defined populations of striatal neurons (Pollak Dorocic *et al*. 2014; Ährlund-Richter *et al*. 2019; Kondabolu *et al*. 2023). We first aimed to characterize and quantify the pallidal neurons that constitute the pallidostriatal projection *as a whole*. The majority of striatal cells are GABAergic, apart from cholinergic interneurons (Tepper *et al*. 2010, 2018; Silberberg and Bolam 2015). In order to selectively target GABAergic neurons, we utilized expression of the *Slc32a1* gene which encodes the vesicular GABA transporter (VGAT), and cholinergic neurons were targeted by the expression of the choline acetyltransferase (ChAT) gene. Accordingly, we unilaterally injected the Cre-dependent “helper virus” (bicistronically expressing TVA receptor fused to a V5 tag, and the rabies glycoprotein [RG] (Ährlund-Richter *et al*. 2019; Kondabolu *et al*. 2023) into the centre of dorsal striatum in double transgenic adult VGAT-Cre:ChAT-Cre mice (n=4). After allowing time for Cre-mediated recombination, and the generation of ‘starter’ neurons expressing all of the proteins necessary for retrograde labelling of pre-synaptic neurons, we injected the modified rabies virus (RG deleted, EnvA, and expressing enhanced GFP) in the same aspect of the striatum (Fig. 1A). Once the modified rabies has replicated and travelled retrogradely, further replication is inhibited due to deletion of endogenous RG (Ährlund-Richter *et al*. 2019).

**Figure 1.**
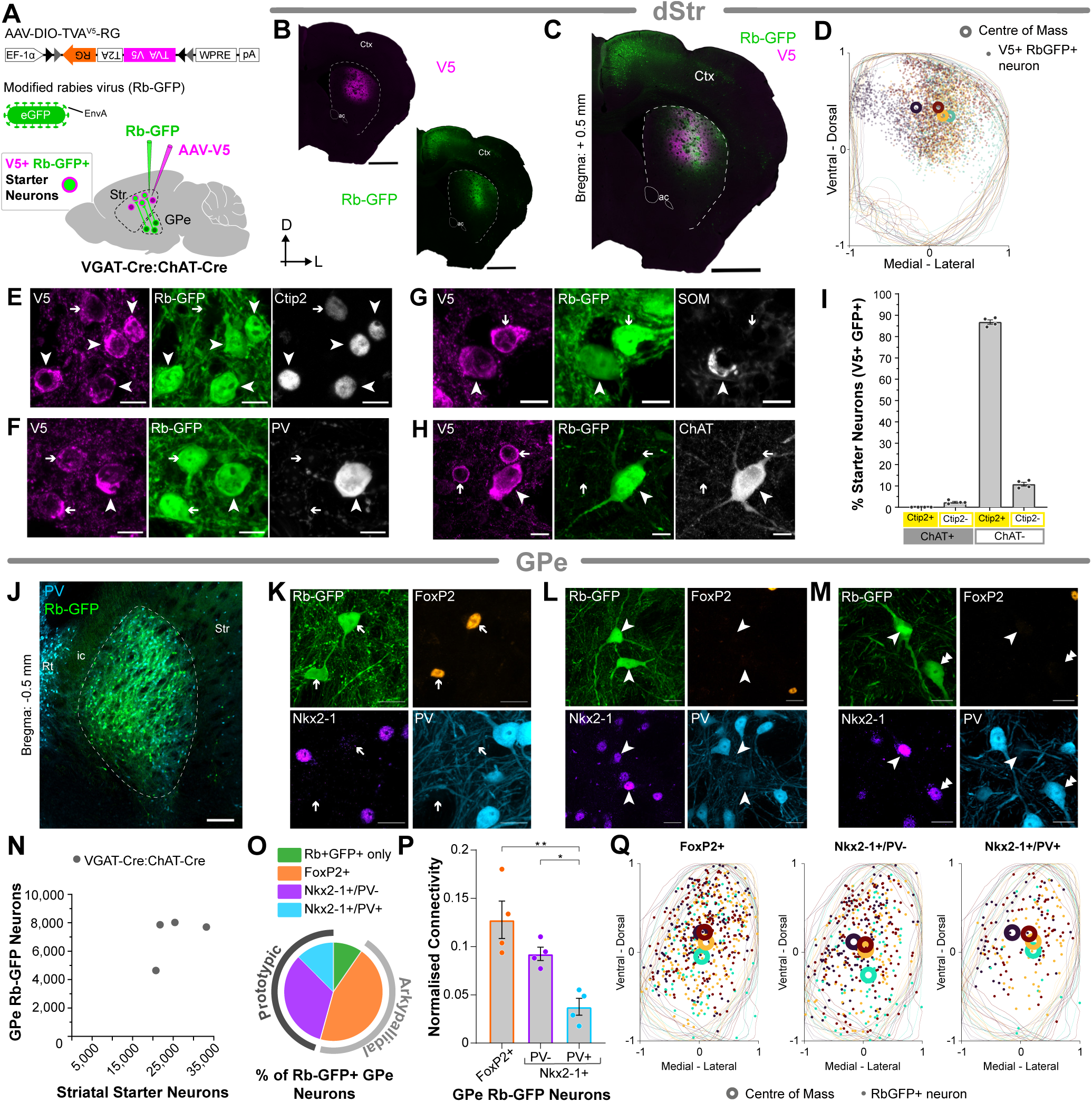
Quantification of neuronal cell types constituting the pallidostriatal pathway. ***A***, Strategy for transsynaptic retrograde labelling of GPe neurons that innervate dorsal striatum. A ‘helper virus’ (AAV-DIO-TVA^V5^-RG) and a modified rabies virus (RG deleted, pseudotyped with EnvA, and expressing eGFP) were unilaterally and sequentially injected into the dorsal striatum of VGAT-Cre:ChAT-Cre mice. Striatal starter neurons, from which transsynaptic labelling of input neurons emanates, co-express V5 (magenta) and rabies-encoded enhanced GFP (Rb-GFP, green). Retrogradely-labelled neurons in GPe (and other brain regions) that innervate the starter neurons express Rb-GFP, but not V5. ***B-C***, Immunofluorescence signals for V5 (***B***, *left*), for Rb-GFP, as expressed by neurons transduced by the rabies virus (***B***, *right*), or for both V5 and Rb-GFP (***C***), in forebrain sections from a single VGAT-Cre:ChAT-Cre mouse. ac, anterior commissure; Ctx, cortex; D, dorsal; L, lateral. ***D***, Distribution of starter neurons (V5+ Rb-GFP+; dots) across multiple coronal sections of the striatum. Each dot represents a neuron counted within the 10 µm optical plane of each coronal section. Colours are per animal (n=4). The mean centre (centre of mass) was calculated for each animal (open circles; n=4). ***E-H***, Immunofluorescence for V5, Rb-GFP and proteins expressed by striatal neuron subtypes in VGAT-Cre:ChAT-Cre mice. ***E***, Neurons expressing both V5 and Rb-GFP (starter neurons; V5+ Rb-GFP+) can also express Ctip2, a marker of SPNs (arrowheads). Note that not all neurons positive for V5 are also positive for Rb-GFP (arrow). ***F***, Some starter neurons were also found to express PV (arrowhead). Note the V5+ Rb-GFP+ neurons that were negative for PV, putative SPNs (arrows). ***G***, Some starter neurons expressed SOM (arrowhead); note the V5+ Rb-GFP+ neuron that is negative for SOM, putative SPN (arrow). ***H***, A subset of starter neurons expressed ChAT (arrowhead); note the smaller V5+ neurons that do no co-express ChAT (putative SPNs; arrows). ***I***, The proportion of striatal starter neurons (V5+ Rb-GFP+) co-expressing Ctip2 or ChAT, as markers of SPNs or cholinergic interneurons, respectively. Unbiased stereology was used to quantify protein co-expression in the striatum of VGAT-Cre:ChAT-Cre mice (n=4). ***J***, Retrogradely labelled Rb-GFP neurons were transduced throughout the GPe, delineated based on the expression of PV (light blue; dashed line indicates boundary). ic, internal capsule; Rt, reticular thalamus; Str, striatum. ***K-L***, Immunofluorescence for markers of pallidal neuronal subtypes in retrogradely labelled neurons. Examples of Rb-GFP neurons (green) in the GPe co-expressing either FoxP2 (***K***, orange; arrows), or Nkx2-1 alone (***L***, purple; arrowheads) or Nkx2-1 and PV (***M***, light blue, stacked arrows). ***N***, Numbers of striatal starter neurons and retrogradely-labelled GPe neurons in VGAT-Cre:ChAT-Cre mice (*n* = 4), as estimated using unbiased stereology (each grey dot indicates estimates from a single mouse). ***O***, Proportion of retrogradely labelled GPe (Rb-GFP+) neurons co-expressing each of the markers assessed. ***P***, Stereological estimates of the subtypes of retrogradely labelled Rb-GFP+ GPe neurons that comprise the pallidostriatal pathway. Normalized connectivity for GPe cells expressing FoxP2 or Nkx2-1 with or without co-expression of PV. Asterisks denote *P < 0.05, and **P < 0.01; one-way ANOVA with Tukey’s post-hoc test. ***Q***, Distribution of Rb-GFP+ neurons (dots) across multiple coronal sections of the GPe, shown for each of the cell types examined: FoxP2+ (*left*), Nkx2-1+/PV- (*middle*) and Nkx2-1+/PV+ (*right*). Each dot represents a neuron counted within the 10 µm optical plane of each coronal section. Colours are per animal (n=4). The mean centre (centre of mass) was calculated for each of the cell types examine, for each animal (open circles). Scale bars: ***B***-***C***, 1 mm; ***E-H***, 10 µm; ***J***, 200 µm; ***K-M***, 20 µm.

We visualized starter neurons in the striatum by examining immunoreactivity for both V5 and the rabies encoded GFP (Rb-GFP; Fig 1B-C). The estimated total number of starter neurons, expressing both V5 and Rb-GFP, was determined using unbiased design-based stereology (Table S2, S3). To ensure that starter neurons were restricted to the central striatum, the position (*x*, *y*) of each counted neuron, within each section, was identified and visualized (Fig 1D; coloured per animal; n=4). To do this, the co-ordinates for the contours and for each counted neuron were exported and placed within a normalized framework (−1 to 1 ventral to dorsal, and −1 to 1 medial to lateral; Fig 1D). This allowed the mean centre or ‘centre of mass’ to be calculated and compared across animals, (open circles; n=4; Fig 1D).

To assess the selectivity of our targeting approach, and to ensure we were capturing SPNs, we carried out immunohistochemistry for Ctip2 as a selective marker of SPNs (Arlotta *et al*. 2008; Garas *et al*. 2016, 2018) (Fig 1E). To ensure that we were also capturing the main populations of GABAergic interneurons we tested immunoreactivity for parvalbumin (PV; Fig 1F) and somatostatin (SOM; Fig 1G). To ensure that we were also capturing cholinergic interneurons we tested immunoreactivity for ChAT (Fig 1H). To rigorously assess the number of starter cells and their phenotype we used unbiased stereology to estimate the proportion of Ctip2 and/or ChAT expressing starter neurons (Fig 1I, Table S3). The majority of neurons expressed Ctip2 but not ChAT (86.8% ± 1.0 %; n=4) typical of SPNs, some neurons expressed ChAT but not Ctip2, thus cholinergic interneurons (2.4% ± 0.4%; n=4) and some neurons expressed neither Ctip2 nor ChAT (10.8% ± 0.9%; n=4) representing a population of putative GABAergic interneurons.

Retrogradely-labelled neurons that monosynaptically innervate starter neurons were identified by their expression of Rb-GFP, but not V5, and were visualized throughout the GPe (Fig 1J). To establish which subsets of pallidal neurons project to the striatum as a whole, we performed immunohistochemistry for known markers of pallidal cell types, FoxP2 as a marker of arkypallidal cells and Nkx2-1 as a marker of prototypic pallidal cells as well as PV as a marker of a sub-set of prototypic neurons. In the VGAT-Cre:ChAT-Cre mice, Rb-GFP+ neurons were found to co-express FoxP2, indicative of arkypallidal neurons (Fig 1K). Rb-GFP+ neurons could also co-express Nkx2-1, either without the co-expression of PV (Nkx2-1+/PV-; Fig 1L) or with PV (Nkx2-1+/PV+; Fig 1M), thus indicative of both sub-types of prototypic neurons. These data confirm that both prototypic and arkypallidal neurons project monosynaptically to striatal neurons.

Using design-based stereological sampling methods, we estimated the total numbers of striatal starter neurons and Rb-GFP+ GPe neurons in each VGAT-Cre:ChAT-Cre mouse (Fig 1N). The average estimated total number of starter neurons per mouse was estimated to be 25,275 (± 2,820; n=4; Fig 1N, Table S3) whereas the average estimated total number of Rb-GFP+ GPe neurons was 7,055 (± 807.65; n=4; Fig 1N, Table S4). To evaluate the relative connectivity from these stereological estimates, we calculated a ‘*normalized connectivity index*’ for each mouse; whereby the number of retrogradely labelled GPe neurons are normalized to the number of striatal starter neurons, this accounts for variation in starter neuron numbers across animals and genotypes (Do *et al*. 2016; Choi *et al*. 2019). The average overall connectivity index for the pallidostriatal pathway in the VGAT-Cre:ChAT-Cre mice was 0.28 (± 0.03; n=4); which indicates, that on average, 100 striatal neurons receive synaptic input from 28 GPe neurons.

In order to gauge the relative contributions of distinct pallidal cell types we evaluated the normalized connectivity for each population of GPe neurons (Fig 1O,P). Despite the fact that FoxP2+ neurons only make up ∼20-30% of GPe cells overall (Abdi *et al*. 2015; Nambu and Chiken 2024) they are the major striatal-projecting single cell type, accounting for 44.7% (± 1.8%; n=4) of retrogradely labelled Rb-GFP+ cells (Fig 1O). Nkx2-1-positive cells, not expressing PV, make up 33.4% (± 2.7%; n=4, Fig 1O), and neurons co-expressing both Nkx2-1 and PV contribute 12.8% (± 1.8%; n=4, Fig 1O). A proportion of retrogradely labelled neurons not expressing any of the markers that we tested were also labelled (Rb-GFP only; 9.1% ± 0.8%; n=4, Fig 1O), as their specific cell type is unknown, we did not include them in any further analyses.

These data demonstrate that the pallidostriatal pathway is composed of neurons from each of the major subsets of pallidal neuronal populations. Furthermore, direct comparisons of the normalized connectivity of each cell type showed that FoxP2 neurons are the major contributors to the pallidostriatal pathway, and that both FoxP2+ and Nkx2-1+/PV-neurons are significantly more connected with the striatum than PV+/Nkx2-1+ neurons (Fig 1P; FoxP2 p=0.0021; Nkx2-1+/PV-p=0.0357; one-way ANOVA with Tukey’s post hoc test; n=4).

The position (*x*, *y*) of each retrogradely labelled Rb-GFP+ counted neuron in the GPe, within the optical disector in each section, was identified and visualized for each animal (Fig 1Q; n=4; coloured per animal). The ‘centre of mass’ for each cell type was then calculated (open circles; Fig 1Q; n=4). Similar to the striatum (Fig 1D) the averaged centre for each animal, and for each cell type, was located centrally within the GPe, and did not show any substantial bias in the medial-lateral or ventral-dorsal aspects. This provides evidence that any comparisons between pallidal cell types innervating the striatum are valid and not due to an underlying difference in the topography of injections, as determined by the location of starter neurons.

### Selective Innervation of Direct Pathway SPNs

Once we evaluated the overall projection to the striatum from the GPe, and the relative contributions of distinct GPe cell populations, we then focused on the pallidal innervation of the major striatal cell types individually. For the objective of this study, we concentrated on the two main classes of projection neurons, the direct and indirect pathway SPNs; to target these populations we used the Drd1a-Cre and Adora2a-Cre mouse lines, respectively (Gerfen *et al*. 1990; Schiffmann and Vanderhaeghen 1993). Direct pathway SPNs (dSPNs) project downstream to the output nuclei of the basal ganglia (Smith *et al*. 1998), however, a proportion of these neurons also have collaterals in the GPe (Kawaguchi, Wilson and Emson 1990; Wu, Richard and Parent 2000; Lévesque and Parent 2005; Fujiyama *et al*. 2011). There is evidence that GPe cells synaptically target striatal projection neurons (Smith *et al*. 1998; Mallet *et al*. 2012), and more specifically dSPNs (Guo *et al*. 2015) however rigorous quantification of the sub-types of neurons in the GPe forming these connections has not yet been undertaken.

In order to examine the nature of pallidal innervation of dSPNs we injected the helper virus and the modified rabies virus into the central striatum of Drd1a-Cre mice (Fig 2A-C; n=5). Neurons expressing both V5 and Rb-GFP (starter neurons) were rigorously counted using unbiased stereology (Table S3) and the location of each counted neuron was identified and visualized to ensure that the starter neurons were restricted to the central dorsal striatum (Fig 2D; coloured per animal; n=5) To evaluate the restriction of the helper virus (V5) and Rb-GFP to dSPNs we labelled neurons for Ctip2, as a marker of all SPNs, and preproenkepahlin (PPE), a marker of indirect pathway SPNs (iSPNs) (Lee *et al*. 1997; Gerfen and Surmeier 2011; Sharott *et al*. 2017) (Fig 2E).

**Figure 2.**
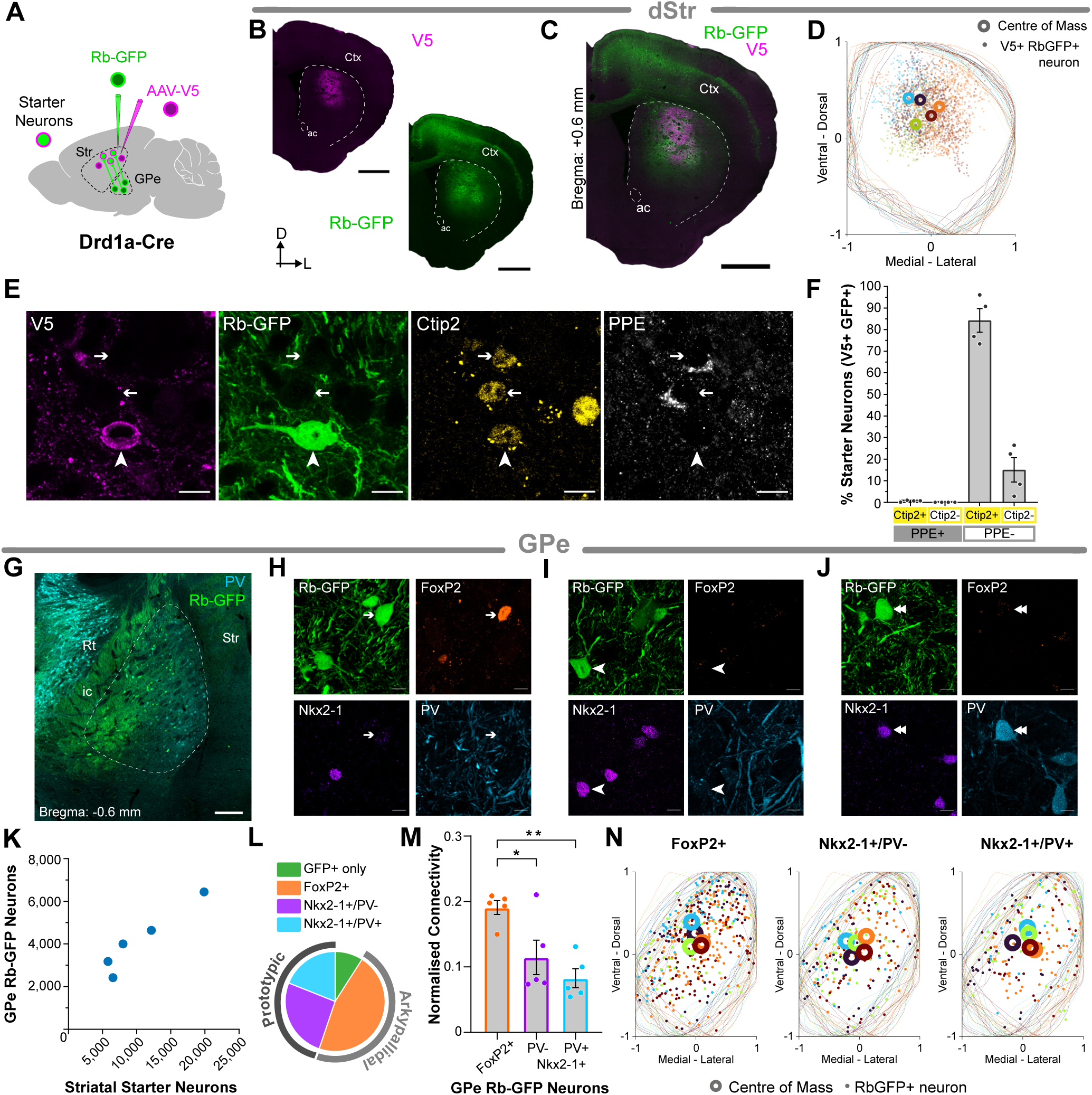
Pallidal innervation of dSPNs. ***A***, Strategy for transsynaptic retrograde labelling of GPe neurons that innervate dSPNs in the dorsal striatum. A ‘helper virus’ (AAV-V5) and a modified rabies virus (Rb-GFP) were unilaterally and sequentially injected into the dorsal striatum of Drd1a-Cre mice. Striatal starter neurons co-express V5 (magenta) and rabies-encoded enhanced GFP (Rb-GFP, green). Retrogradely-labelled neurons in GPe (and other brain regions) that innervate the starter neurons express Rb-GFP, but not V5. ***B-C***, Immunofluorescence signals for V5 (***B***, *left*), for Rb-GFP, as expressed by neurons transduced by the rabies virus (***B,*** *right*), or for both V5 and Rb-GFP (***C***), in forebrain sections from a single Drd1a-Cre mouse. ac, anterior commissure; Ctx, cortex; D, dorsal; L, lateral. ***D***, Distribution of V5+ Rb-GFP+ starter neurons (dots) across multiple coronal sections of the striatum. Each dot represents a neuron counted within the 10 µm optical disector of each section. Colours are per animal. The mean centre (centre of mass) was calculated for each animal (open circles; n=5). ***E***, Immunofluorescence for V5, Rb-GFP and proteins expressed by striatal neuron subtypes in Drd1a-Cre mice. Neurons expressing both V5 and Rb-GFP (starter neurons; V5+/Rb-GFP+) can also express Ctip2, a marker of SPNs (arrowhead). Note that these neurons are not positive for PPE, a marker of iSPNs. Note that some Ctip2 labelled cells also express PPE and are negative for both V5 and Rb-GFP, likely iSPNs (arrows). ***F***, The proportion of striatal starter neurons (V5+/Rb-GFP+) co-expressing Ctip2 and/or PPE, as markers of all SPNs or iSPNs, respectively. Unbiased stereology was used to quantify protein co-expression in the striatum of Drd1a-Cre mice (n=4). ***G***, Retrogradely labelled Rb-GFP neurons were transduced in the GPe, delineated based on the expression of PV (light blue; dashed line indicates boundary). ic, internal capsule; Rt, reticular thalamus; Str, striatum. ***H-J***, Immunofluorescence for markers of pallidal neuronal subtypes in retrogradely labelled neurons. Examples of Rb-GFP neurons in the GPe co-expressing FoxP2 (***H***, orange; arrow), Nkx2-1+/PV- (***I***, purple; arrowhead) or Nkx2-1+/PV+ (***J***, purple, light-blue, stacked arrowhead). ***K***, Numbers of striatal starter neurons and retrogradely-labelled Rb-GFP+ GPe neurons in Drd1a-Cre mice (*n* = 5), as estimated using design-based stereology (each blue dot indicates estimates from a single mouse). ***L***, Proportions (%) of retrogradely labelled GPe (Rb-GFP+) neurons co-expressing each of the markers tested. ***M***, Stereological estimates of the subtypes of retrogradely labelled Rb-GFP+ GPe neurons that comprise the pallidostriatal pathway innervating dSPNs. Normalised connectivity for GPe cells expressing FoxP2 or Nkx2-1 with or without co-expression of PV. Asterisks denote *P < 0.05, and **P < 0.01; one-way ANOVA with Tukey’s post-hoc test. ***N***, Distribution of Rb-GFP+ neurons (dots) across multiple coronal sections of the GPe, shown for each of the cell types examined: FoxP2+ (*left*), Nkx2-1+/PV- (*middle*) and Nkx2-1+/PV+ (*right*). Each dot represents a neuron counted within the 10 µm optical disector of each section. Colours are per animal. The mean centre (centre of mass) was calculated for each of the cell types examined, for each animal (open circles; n=5). Scale bars: ***B***-***C***, 1 mm; ***E***, 10 µm; ***G***, 200 µm; ***H-J***, 10 µm.

Stereological quantification of starter cells showed that the majority expressed Ctip2 and not PPE (84.5% ± 5.5% n =4; Fig 2F), typical of dSPNs. A proportion of starter cells did not express Ctip2, nor PPE (15.0% ± 5.6%; n=4; Fig 2F), and less than 1% of neurons expressed PPE with or without the presence of Ctip2 (Fig. 2F). This indicates that the majority of starter cells have a phenotype consistent with dSPNs.

Retrogradely labelled Rb-GFP neurons were visualized within the GPe (Fig. 2G). To establish which subsets of pallidal neurons innervate dSPNs, we performed immunohistochemistry for FoxP2, Nkx2-1 and PV (Fig 2H-J). Retrogradely labelled Rb-GFP GPe neurons were shown to co-express all the markers that we tested, demonstrating that both prototypic and arkypallidal neurons innervated dSPNs (Fig 2H-J).

Using stereology, we estimated the total numbers of striatal starter neurons and Rb-GFP+ GPe neurons in each Drd1a-Cre mouse (Fig 2K). The average estimated total number of starter neurons per mouse was 10,470 (± 2,598; n=5) and the average estimated total number of Rb-GFP+ GPe neurons was 4,136 (± 687; n=5). The average overall connectivity index in the Drd1a-Cre mice was 0.43 (± 0.04; n=5) GPe Rb-GFP cells per starter cell.

We then looked at the relative contribution of distinct pallidal cell types projecting to dSPNs (Fig 2L). The major single cell type is FoxP2+ neurons, constituting 46.1% (± 3.99%; n=5); followed by Nkx2-1+/PV-neurons making up 25.8% (± 3.33%; n=5); then Nkx2-1+/PV+ neurons with 19.0% (± 1.56%; n=5); and GFP only neurons (9.06% ± 2.12%; n=5; Fig 2L). We evaluated the normalized connectivity for each population of GPe neurons projecting to dSPNs (Fig 2M). Arkypallidal FoxP2+ neurons project to dSPNs significantly more compared to Nkx2.1+/PV- and Nkx2-1+/PV+ neurons (p=0.0314 and p=0.0035, respectively; one-way ANOVA with Tukey’s post hoc test; n=5; Fig 2M).

The position (*x*, *y*) of each retrogradely labelled Rb-GFP neuron in the GPe, within each section, was identified and visualized (Fig 2N; n=5; coloured per animal). The ‘centre of mass’ for each cell type was then calculated (Fig 2N). Similar to the striatum (Fig 2D), the centre of all counted Rb-GFP+ neurons, for each cell type, in each animal, did not show any major bias in either the ventral-dorsal or medial-lateral axes.

### Selective Innervation of Indirect Pathway SPNs

Indirect pathway SPNs (iSPNs) exert their influence over the output of the basal ganglia via the GPe; thus, understanding the reciprocal connectivity between GPe neurons and iSPNs will undoubtedly help clarify the underlying function of this network. As stated previously, there is evidence that GPe neurons target SPNs, and there is a described projection directly to iSPNs (Guo *et al*. 2015); however, the exact subsets of GPe neurons projecting to this population remain unknown.

To study the innervation of iSPNs we injected the helper and the modified rabies viruses into the striatum of Adora2a-Cre mice (Fig 3A-C). The total number of starter neurons, expressing both V5 and Rb-GFP, was quantified using stereology (Table S3). The co- ordinates for each starter neuron (*x*, *y*) were identified and visualized; and the centre of mass of the starter neurons per animal was calculated (Fig 3D).

**Figure 3.**
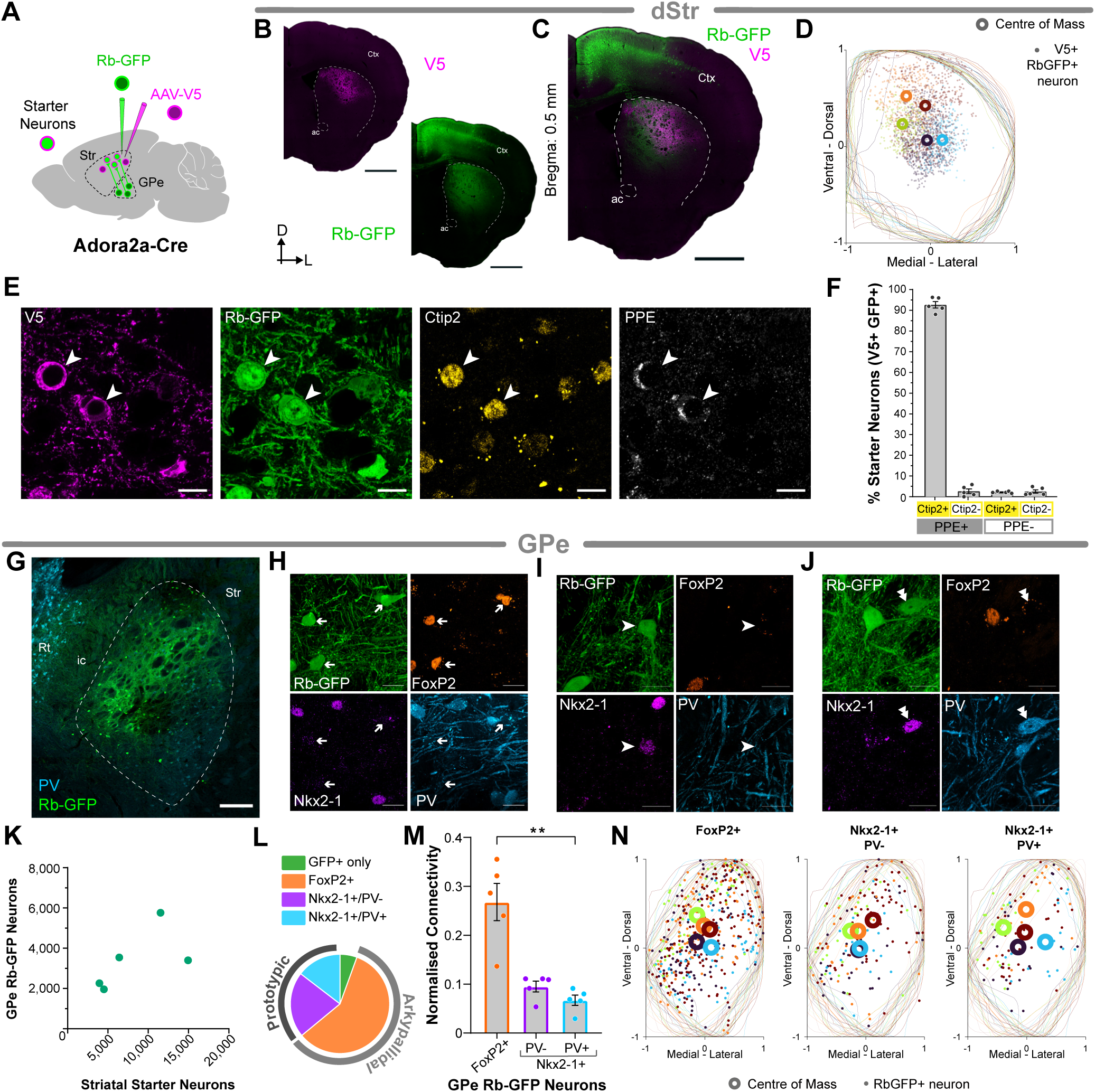
Pallidal innervation of iSPNs. ***A***, Strategy for transsynaptic retrograde labelling of GPe neurons that innervate iSPNs in the dorsal striatum. A ‘helper virus’ (AAV-V5) and a modified rabies virus (Rb-GFP) were unilaterally and sequentially injected into the dorsal striatum of Adora2a-Cre mice. Striatal starter neurons co-express V5 (magenta) and rabies-encoded enhanced GFP (Rb-GFP, green). Retrogradely-labelled neurons in GPe (and other brain regions) that innervate the starter neurons express Rb-GFP, but not V5. ***B-C***, Immunofluorescence signals for V5 (***B****, left*), for Rb-GFP, as expressed by neurons transduced by the rabies virus (***B***, *right*), or for both V5 and Rb-GFP (***C***), in forebrain sections from a single Adora2a-Cre mouse. ac, anterior commissure; Ctx, cortex; D, dorsal; L, lateral. ***D***, Distribution of V5+ Rb-GFP+ starter neurons (dots) across multiple coronal sections of the striatum. Each dot represents a neuron counted within the 10 µm optical plane of each section. Colours are per animal. The mean centre (centre of mass) was calculated for each animal (open circles; n=5). ***E***, Immunofluorescence for V5, Rb-GFP and proteins expressed by striatal neuron subtypes in Adora2a-Cre mice. Neurons expressing both V5 and Rb-GFP (starter neurons; V5+ Rb-GFP+) can also express Ctip2 (yellow) and PPE (white; arrowheads), markers of all SPNs, and iSPNs, respectively. ***F***, The proportion of striatal starter neurons (V5+/Rb-GFP+) co-expressing Ctip2 and PPE as markers of iSPNs. Design-based stereology was used to quantify protein co-expression in the striatum of Adora2a-Cre mice (n=5). ***G***, Retrogradely labelled Rb-GFP neurons were transduced in the GPe, delineated based on the expression of PV (light blue; dashed line indicates boundary). ic, internal capsule; Rt, reticular thalamus; Str, striatum. ***H-J***, Immunofluorescence for markers of pallidal neuronal subtypes in retrogradely labelled neurons. Examples of Rb-GFP neurons in the GPe co-expressing FoxP2 (***H***, orange; arrow), Nkx2-1+/PV- (***I***, purple; arrowhead) or Nkx2-1+/PV+ (***J***, purple, light-blue, stacked arrowhead). ***K***, Numbers of striatal starter neurons and retrogradely-labelled Rb-GFP+ GPe neurons in Adora2a-Cre mice (*n* = 5), as estimated using design-based stereology (each green dot indicates estimates from a single mouse). ***L***, Proportions (%) of retrogradely labelled GPe (Rb-GFP+) neurons co-expressing each of the markers tested. ***M***, Stereological estimates of the subtypes of retrogradely labelled Rb-GFP+ GPe neurons that comprise the pallidostriatal pathway innervating iSPNs. Normalised connectivity for GPe cells expressing FoxP2 or Nkx2-1 with or without co-expression of PV. Asterisks denote **P < 0.01; Kruskal-Wallis with Dunn’s post-hoc test. ***N***, Distribution of Rb-GFP+ neurons (dots) across multiple coronal sections of the GPe, shown for each of the cell types examined: FoxP2+ (*left*), Nkx2-1+ PV- (*middle*) and Nkx2-1+ PV+ (*right*). Each dot represents a neuron counted within the 10 µm optical plane of each section. Colours are per animal. The mean centre (centre of mass) was calculated for each of the cell types examine, for each animal (open circles; n=5). Scale bars: ***B***-***C***, 1 mm; ***E***, 10 µm; ***G***, 200 µm; ***H-J***, 20 µm.

To confirm selectivity of the tracing strategy we tested immunoreactivity for V5, Rb-GFP, Ctip2 and PPE (Fig 3E). Quantitative analyses with stereology showed that most starter neurons expressed both Ctip2 and PPE (92.6% ± 1.6%, n=5; Fig 3F). A smaller proportion expressed PPE but not Ctip2 (2.7% ± 1.1%, n=5), or neither Ctip2 nor PPE (2.5% ± 0.8%, n=5) or Ctip2 and not PPE (2.20% ± 0.20%, n = 5; Fig 2G). This data confirms that most starter neurons have a phenotype consistent with iSPNs.

Analysis of the GPe confirmed the presence of retrogradely labelled Rb-GFP neurons throughout, confirming a monosynaptic pallidostriatal projection from the GPe to iSPNs (Fig 3G). Further analysis revealed that Rb-GFP+ neurons co-express known markers of GPe neuron subtypes, FoxP2 or Nkx2-1 with and without PV (Fig 3H-J). The data thus demonstrates that both arkypallidal and prototypic neurons are synaptically connected with iSPNs.

Using stereology, we estimated the total numbers of striatal starter neurons and Rb-GFP+ GPe neurons in each Adora2a-Cre mouse (Fig 3K). The average estimated total number of starter neurons per mouse was 8,263 (± 2,139; n=5;) and the average estimated total number of Rb-GFP+ GPe neurons was 3,384 (± 669; n=5). The average overall connectivity index in the Adora2a-Cre mice was 0.46 (± 0.06; n=5) GPe Rb-GFP cells per starter cell.

Out of all GPe neurons projecting to iSPNs FoxP2+ arkypallidal neurons show the greatest connectivity (58.5% ± 1.2), followed by Nkx2-1+/PV-neurons (21.4% ± 1.7) and then Nkx2-1+/PV+ neurons (14.6% ± 1.1; Fig. 3L). The remaining neurons, only expressing Rb-GFP+, make up 5.5% (±1.5%) of the total population of GPe neurons projecting to iSPNs (Fig. 3L). We evaluated the normalized connectivity for each population of GPe neurons projecting to dSPNs (Fig 3M). Arkypallidal FoxP2+ neurons are significantly more connected with iSPNs than Nkx2-1+/PV+ GPe neurons (p = 0.004, Kruskal-Wallis with Dunn’s; Fig. 3M).

The position (*x*, *y*) of each retrogradely labelled Rb-GFP neuron in the GPe, within each section, was identified and visualized (Fig 3N; n=5; coloured per animal). The ‘centre of mass’ for each cell type was then calculated (Fig 3N). Similar to the striatum (Fig 3D), the centre of Rb-GFP neurons, for each cell type, in each animal, did not show any major bias in either the ventral-dorsal or medial-lateral planes.

### Selective Innervation of Striatal Parvalbumin-Expressing Interneurons

Aside from the vastly numerous SPNs, there are at least two major GABAergic interneuron populations: those expressing either parvalbumin (PV) or somatostatin (SOM) (Kawaguchi *et al*. 1995; Sharott *et al*. 2012). Despite the relatively low numbers of interneurons, studies have continued to show that these neurons are in a key position to alter striatal dynamics and have selective connectivity with other brain regions (Straub *et al*. 2016; Johansson and Silberberg 2020). Thus, elucidating the specific pallidal inputs to interneurons will provide important structural features that no doubt has important functional correlates.

A projection from the GPe to striatal PV interneurons has previously been described (Klug *et al*. 2018) but to precisely define and quantify the population of pallidal neurons that give rise innervation, the helper and rabies viruses were sequentially injected into the dorsal aspects of the striatum of PV-Cre mice (Fig 4A-C). The total number of starter neurons, expressing both V5 and Rb-GFP, was quantified using stereology (Table S3). The co-ordinates for each starter neuron (*x*, *y*) were identified and visualized; and the centre of mass for the starter neurons per animal was calculated (Fig 4D; n=9).

**Figure 4.**
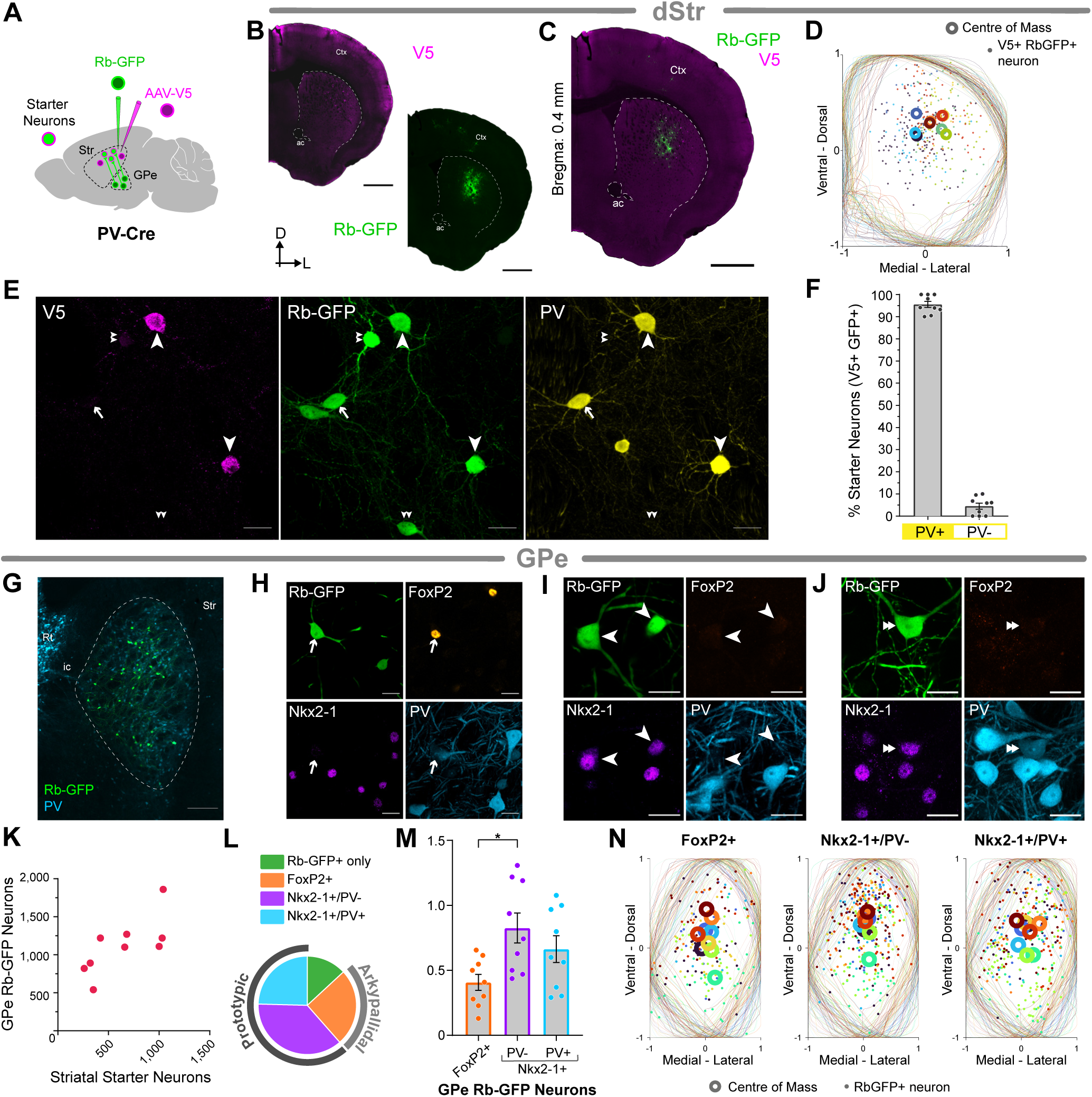
Pallidal innervation of striatal PV interneurons. ***A***, Strategy for transsynaptic retrograde labelling of GPe neurons that innervate PV interneurons in the dorsal striatum. A ‘helper virus’ (AAV-V5) and a modified rabies virus (Rb-GFP) were unilaterally and sequentially injected into the dorsal striatum of PV-Cre mice. Striatal starter neurons co-express V5 (magenta) and rabies-encoded enhanced GFP (Rb-GFP, green). Retrogradely-labelled neurons in GPe (and other brain regions) that innervate the starter neurons express Rb-GFP, but not V5. ***B-C***, Immunofluorescence signals for V5 (***B***, *left*), for Rb-GFP, as expressed by neurons transduced by the rabies virus (***B,*** *right*), or for both V5 and Rb-GFP (***C***), in forebrain sections from a single PV-Cre mouse. ac, anterior commissure; Ctx, cortex; D, dorsal; L, lateral. ***D***, Distribution of V5+/Rb-GFP+ starter neurons (dots) across multiple coronal sections of the striatum. Each dot represents a neuron counted within the 10 µm optical plane of each section. Colours are per animal. The mean centre (centre of mass) was calculated for each animal (open circles; n=9). ***E***, Immunofluorescence for V5, Rb-GFP and parvalbumin (PV). Neurons expressing both V5 and Rb-GFP (starter neurons; V5+/Rb-GFP+) can also express PV (arrowheads). Note that some Rb-GFP+ neurons did not express V5 or PV (putative SPNs; double arrowheads) and that not all PV+ neurons were positive for V5 and Rb-GFP (arrow). ***F***, The proportion of striatal starter neurons (V5+/Rb-GFP+) co-expressing PV was determined using design-based stereology in PV-Cre mice (n=9). ***G***, Retrogradely labelled Rb-GFP neurons were transduced in the GPe, delineated based on the expression of PV (light blue; dashed line indicates boundary). ic, internal capsule; Rt, reticular thalamus; Str, striatum. ***H-J***, Immunofluorescence for markers of pallidal neuronal subtypes in retrogradely labelled neurons. Examples of Rb-GFP neurons in the GPe co-expressing FoxP2 (***H***, orange; arrow) or Nkx2-1 (***I***, purple; arrowheads) or Nkx2-1 with PV (***J***, light blue, stacked arrowheads). ***K***, Total numbers of striatal starter neurons and retrogradely-labelled Rb-GFP+ GPe neurons in PV-Cre mice (*n* = 9), as estimated using design-based stereology (each pink dot indicates estimates from a single mouse). ***L***, Proportion of retrogradely labelled GPe (Rb-GFP+) neurons co-expressing each of the markers evaluated. ***M***, Stereological estimates of the subtypes of retrogradely labelled Rb-GFP+ GPe neurons that comprise the pallidostriatal pathway innervating PV interneurons. Normalised connectivity for GPe cells expressing FoxP2 or Nkx2-1 with or without co-expression of PV. Asterisk denotes *P < 0.05, one-way ANOVA with Tukey’s post-hoc test. ***N***, Distribution of Rb-GFP+ neurons (dots) across multiple coronal sections of the GPe, shown for each of the cell types examined: FoxP2+ (*left*), Nkx2-1+ PV- (*middle*) and Nkx2-1+ PV+ (*right*). Each dot represents a neuron counted within the 10 µm optical plane of each section. Colours are per animal. The mean centre (centre of mass) was calculated for each of the cell types examined, for each animal (open circles; n=9). Scale bars: ***B***-***C***, 1 mm; ***E***, 20 µm; ***G***, 200 µm; ***H-J***, 20 µm.

To confirm that our tracing strategy was selective for PV-interneurons we examined immunoreactivity for parvalbumin in V5+/Rb-GFP+ starter neurons (Fig 4E). Using stereology, we evaluated the proportion of PV+ expressing starter cells to be 95.5% (±1.4%; n=9) and the remainder were negative for PV (Fig 4F; 4.5% ±1.4%; n=9). Thus, confirming that our tracing strategy was indeed highly selective for PV-interneurons.

Examination of the GPe showed that there were retrogradely labelled Rb-GFP neurons present, confirming a direct projection from the GPe to striatal PV interneurons. Investigation into the sub-types of GPe neurons projecting to striatal PV interneurons revealed that there were Rb-GFP+ neurons also expressing FoxP2, presenting evidence that arkypallidal neurons innervate PV interneurons (Fig 4H). Examination of prototypic markers showed that Rb-GFP could co-express with Nkx2-1 without the presence of PV (Fig 4I) and with PV (Fig 4J). These data confirm that both arkypallidal and prototypic neurons are synaptically connected to striatal PV interneurons.

Using stereology, we estimated the total numbers of striatal starter neurons and Rb-GFP+ GPe neurons in each PV-Cre mouse (Fig 4K). The average estimated total number of starter neurons per mouse was 640.0 (±106.9; n=9; Table S3) and the average estimated total number of Rb-GFP+ GPe neurons was 1,177.6 (± 145.9; n=9; Table S4). The average overall connectivity index in the PV-Cre mice was 2.09 (± 0.26; n=9) GPe Rb-GFP cells per starter cell.

Analysis of the retrogradely labelled pallidal cells projecting to PV interneurons revealed that, unlike the SPNs, the two major cell types innervating PV interneurons are Nkx2-1+ neurons not expressing PV (38.0% ± 1.8; n=9; Fig 4L) and Nkx2-1+ neurons expressing PV (32.9% ± 2.6%; n=9). FoxP2+ Rb-GFP constituted 19.7% (± 1.9%; n=9) of the pallidal cells and Rb-GFP+ only neurons at 9.4% (± 1.3%; n=9; Fig 4L). In addition, Nkx2-1+/PV- neurons are significantly more connected with PV interneurons than FoxP2+ GPe neurons (p = 0.014, One-way ANOVA with Tukey’s post hoc test; Fig. 4M).

The position (*x*, *y*) of each retrogradely labelled Rb-GFP neuron in the GPe, within each section, was identified and visualized (Fig 4N; n=9; coloured per animal). The ‘centre of mass’ for each cell type was then calculated (Fig 4N). Like the striatum (Fig 4D), the centre of Rb-GFP neurons, for each cell type, in each animal, did not show any major bias in either the ventral-dorsal or medial-lateral planes.

### Selective Innervation of Striatal Somatostatin-Expressing Interneurons

Somatostatin (SOM)-expressing interneurons are a subset of striatal GABAergic cells, once thought to be a uniform group co-expressing neuropeptide Y (NPY) and nitric oxide synthase (NOS). In mice, SOM/NOS-positive neurons are classified as PLTS interneurons, distinct from NPY-neurogliaform neurons lacking SOM (Ibanez-Sandoval *et al*. 2010). These subtypes differ in structure and function (English *et al*. 2012; Holly *et al*. 2019). This study focuses on SOM-expressing PLTS neurons, implicated in goal-directed learning (Holly *et al*. 2019). Pallidostriatal inputs target NOS-positive neurons in rats (Bevan *et al*. 1998), and in mice, SOM interneurons receive GPe input (Choi *et al*. 2019), though the exact pallidal sources remain unclear.

To investigate the precise pallidal innervation of these SOM neurons, we injected the helper and rabies viruses into the striatum of SOM-Cre mice (Fig. 5A-C). The total number of starter neurons, expressing both V5 and Rb-GFP, was quantified using stereology (Table S3). To examine the distribution of the starter neurons, the co-ordinates for each starter neuron (*x*, *y*) were exported and placed within a normalized framework (Fig 5D; n=5). The mean centre (centre of mass) of all counted neurons in each section, for each animal, was then calculated (Fig 5D; n=5).

**Figure 5.**
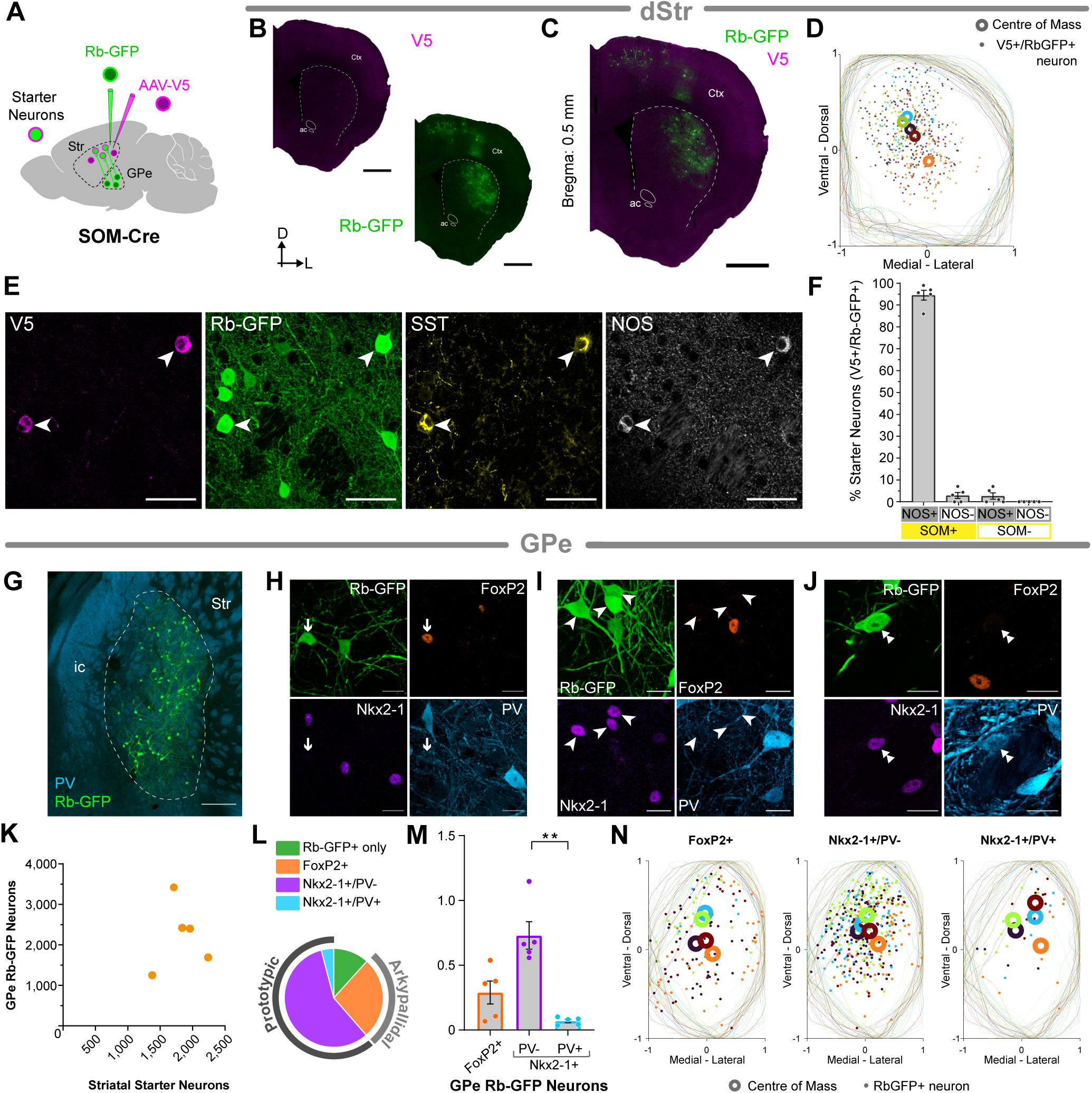
Pallidal innervation of striatal SOM interneurons. ***A***, Strategy for transsynaptic retrograde labelling of GPe neurons that innervate SOM interneurons in the dorsal striatum. A ‘helper virus’ (AAV-V5) and a modified rabies virus (Rb-GFP) were unilaterally and sequentially injected into the dorsal striatum of SOM-Cre mice. Striatal starter neurons co-express V5 (magenta) and rabies-encoded enhanced GFP (Rb-GFP, green). Retrogradely-labelled neurons in GPe (and other brain regions) that innervate the starter neurons express Rb-GFP, but not V5. ***B-C***, Immunofluorescence signals for V5 (***B***, *left*), for Rb-GFP, as expressed by neurons transduced by the rabies virus (***B,*** *right*), or for both V5 and Rb-GFP (***C***), in forebrain sections from a single SOM-Cre mouse. ac, anterior commissure; Ctx, cortex; D, dorsal; L, lateral. ***D***, Distribution of V5+ Rb-GFP+ starter neurons (dots) across multiple coronal sections of the striatum. Each dot represents a neuron counted within the 10 µm optical plane of each section. Colours are per animal. The mean centre (‘centre of mass’) was calculated for each animal (open circles; n=5). ***E***, Immunofluorescence for V5, Rb-GFP, SOM and NOS. Neurons expressing both V5 and Rb-GFP (starter neurons; V5+ Rb-GFP+) can also express SOM and NOS (arrowheads). Note that some Rb-GFP+ neurons did not express V5, SOM or NOS. ***F***, The proportion of striatal starter neurons (V5+ Rb-GFP+) co-expressing SOM and/or NOS was determined using unbiased stereology in SOM-Cre mice (n=5). ***G***, Retrogradely labelled Rb-GFP neurons were transduced in the GPe, delineated based on the expression of PV (light blue; dashed line indicates boundary). ic, internal capsule; Rt, reticular thalamus; Str, striatum. ***H***, Immunofluorescence for markers of pallidal neuronal subtypes in retrogradely labelled neurons. Examples of Rb-GFP neurons in the GPe co-expressing FoxP2 (orange; arrow) or Nkx2-1 (purple; arrowheads). Note that in this example all of the neurons expressing Nkx2-1 do not also express PV, but that there is a positive neuron within the image frame. ***I***, Numbers of striatal starter neurons and retrogradely-labelled Rb-GFP+ GPe neurons in SOM-Cre mice (*n* = 5), as estimated using unbiased stereology (each orange dot indicates estimates from a single mouse). ***J***, Stereological estimates of the subtypes of retrogradely labelled Rb-GFP+ GPe neurons that comprise the pallidostriatal pathway innervating SOM interneurons. Normalised connectivity for GPe cells expressing FoxP2 or Nkx2-1 with or without co-expression of PV. Asterisks denote **P < 0.01, ***P < 0.001; ns, not significant; one-way ANOVA with Tukey’s post-hoc test. ***K***, Distribution of Rb-GFP+ neurons (dots) across multiple coronal sections of the GPe, shown for each of the cell types examined: FoxP2+ (*left*), Nkx2-1+ PV- (*middle*) and Nkx2-1+ PV+ (*right*). Each dot represents a neuron counted within the 10 µm optical plane of each section. Colours are per animal. The mean centre (‘centre of mass’) was calculated for each of the cell types examined, for each animal (open circles; n=5). Scale bars: ***B***-***C***, 1 mm; ***E***, 20 µm; ***G***, 200 µm; ***H***, 10 µm.

To evaluate the specificity of our tracing approach in the SOM-Cre animals, we tested immunoreactivity for SOM and NOS in starter neurons (Fig 5E). We then used unbiased stereology to quantify the co-expression of these proteins (Fig 5F). Most starter neurons expressed both NOS and SOM (94.5 ± 2.2 %; n=5; Fig. 5F). A smaller number of neurons expressed SOM but not NOS (2.9 ± 1.3 %; n=5) or NOS but not SOM (2.6 ± 1.4 %; n=5; Fig. 5F). Thus, the majority of neurons expressed both SOM and NOS and thus have the phenotype of somatostatin-expressing PLTS neurons (Ibáñez-Sandoval *et al*. 2011).

In the GPe of the SOM-Cre injected mice, there were Rb-GFP+ neurons distributed throughout, confirming a direct monosynaptic projection from the GPe to SOM interneurons (Fig 5G). We then analysed the phenotype of these neurons based on co-expression of known GPe cell markers (Fig 5H-J). We found that Rb-GFP+ neurons could co-express FoxP2, indicating that arkypallidal neurons innervate SOM neurons (Fig 5H); and that Rb-GFP could also express markers of prototypic neurons (Fig 5I,J).

Using stereology, we estimated the total numbers of striatal starter neurons and Rb-GFP+ GPe neurons in each SOM-Cre mouse (Fig 5K). The average estimated total number of starter neurons per mouse was 1,825 (±142.3; n=5; Table S3) and the average estimated total number of Rb-GFP+ GPe neurons was 2,236 (±369.9; n=5; Table S4). The average overall connectivity index in the SOM-Cre mice was 1.24 (± 0.22; n=5) GPe Rb-GFP cells per starter cell.

Stereological analysis of the retrogradely labelled Rb-GFP GPe neurons revealed that the main subset of neurons innervating SOM interneurons is prototypic neurons (Nkx2-1+) not expressing PV (61.1% ± 5.6%, n=5; Fig 5L), followed by FoxP2+ neurons (21.3% ± 4.5%; n=5) and then Nkx2.1+/PV+ neurons (5.7% ± 0.8%; n=5). A proportion of neurons were also negative for all the markers that we tested (GFP only; 11.8% ± 1.4%; n=5; Fig 5M). In summary, Nkx2-1 neurons, not expressing PV, are significantly more connected with SOM interneurons than Nkx2.1 neurons co-expressing PV (p= 0.0027; n=5; Kruskal-Wallis with Dunn’s multiple comparisons test; Fig 5M).

The position (*x*, *y*) of each retrogradely labelled Rb-GFP neuron counted within the 10 µm optical disector for every section in each animal was exported so that the data could be visualized in a normalized framework (Fig 5N; n=5; coloured per animal). The ‘centre of mass’ for each cell type was then calculated (Fig 5N). Comparable to the striatum (Fig 5D), the centre of Rb-GFP neurons, for each cell type, in each animal, did not show any major bias in either the ventral-dorsal or medial-lateral planes.

### Selective Innervation of Striatal Cholinergic Interneurons

Aside from GABAergic neurons in the striatum there is also a population of cholinergic interneurons; these large aspiny neurons have a role in associative learning, responding with a ‘conditioned pause’ when a salient stimulus is linked to an aversive or rewarding outcome (Kimura, Rajkowski and Evarts 1984; Aosaki *et al*. 1994; Ravel, Legallet and Apicella 2003; Apicella *et al*. 2011). This pause, and the temporal control of acetylcholine release is intricately related to reward-related learning mediated by dopamine (Wang *et al*. 2006; Apicella *et al*. 2011; Threlfell *et al*. 2012; Nelson *et al*. 2014; Kosillo *et al*. 2016; Zhang, Reynolds and Cragg 2018). Cholinergic interneurons receive innervation from local striatal neurons and a variety of different brain regions (Doig *et al*. 2014; Lim, Kang and McGehee 2014) including the GPe (Guo *et al*. 2015; Choi *et al*. 2019). Comprehensive understanding of the particular neurons underlying the connectivity of cholinergic interneurons will help dissect the physiology underlying the behavioural roles of these interneurons.

To examine the pallidal innervation of cholinergic interneurons, we injected the helper and rabies viruses into the dorsal striatum of ChAT-Cre mice (Fig. 6A-C). The total number of starter neurons, expressing both V5 and Rb-GFP, was quantified using stereology (Table S3). Neurons were counted in a 10 µm optical disector for every section containing V5+ neurons, within a series, per animal. The co-ordinates of these counted neurons were exported and placed within a normalized framework (Fig 6D). The mean centre (centre of mass) for all starter neurons, for each animal was calculated (open circles, Fig 6D; n=5).

**Figure 6.**
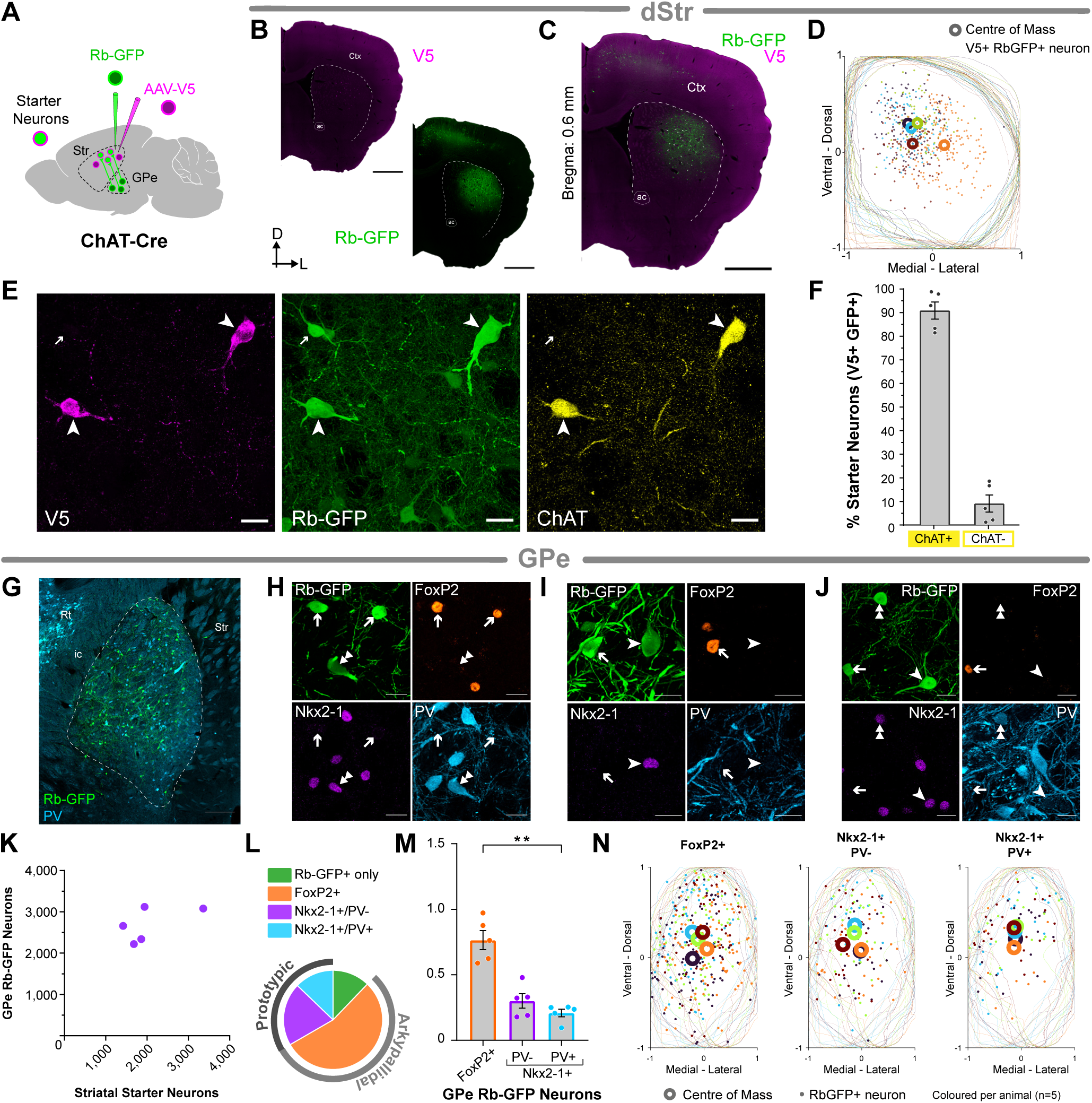
Pallidal innervation of striatal ChAT interneurons. ***A***, Strategy for transsynaptic retrograde labelling of GPe neurons that innervate ChAT interneurons in the dorsal striatum. A ‘helper virus’ (AAV-V5) and a modified rabies virus (Rb-GFP) were unilaterally and sequentially injected into the dorsal striatum of ChAT-Cre mice. Striatal starter neurons co-express V5 (magenta) and rabies-encoded enhanced GFP (Rb-GFP, green). Retrogradely-labelled neurons in GPe (and other brain regions) that innervate the starter neurons express Rb-GFP, but not V5. ***B-C***, Immunofluorescence signals for V5 (***B***, *left*), for Rb-GFP, as expressed by neurons transduced by the rabies virus (***B,*** *right*), or for both V5 and Rb-GFP (***C***), in forebrain sections from a single ChAT-Cre mouse. ac, anterior commissure; Ctx, cortex; D, dorsal; L, lateral. ***D***, Distribution of V5+ Rb-GFP+ starter neurons (dots) across multiple coronal sections of the striatum. Each dot represents a neuron counted within the 10 µm optical plane of each section. Colours are per animal. The mean centre (‘centre of mass’) was calculated for each animal (open circles; n=5). ***E***, Immunofluorescence for V5 (magenta), Rb-GFP (green) and ChAT (yellow). Neurons expressing both V5 and Rb-GFP (starter neurons; V5+ Rb-GFP+) can also express ChAT (arrowheads). Note that some Rb-GFP+ neurons do not express V5 or ChAT (arrow). ***F***, The proportion of striatal starter neurons (V5+ Rb-GFP+) co-expressing SOM and/or NOS was determined using unbiased stereology in SOM-Cre mice (n=5). ***G***, Retrogradely labelled Rb-GFP neurons were transduced in the GPe of ChAT-Cre mice, delineated based on the expression of PV (light blue; dashed line indicates boundary). ic, internal capsule; Rt, reticular thalamus; Str, striatum. ***H***, Immunofluorescence for markers of pallidal neuronal subtypes in retrogradely labelled neurons. Examples of Rb-GFP neurons in the GPe co-expressing FoxP2 (orange; arrows) or Nkx2-1 (purple; arrowhead). Note that, in this example, the neuron shown expressing Nkx2-1 does also express PV. ***I***, Numbers of striatal starter neurons and retrogradely-labelled Rb-GFP+ GPe neurons in ChAT-Cre mice (*n* = 5), as estimated using unbiased stereology (each purple dot indicates estimates from a single mouse). ***J***, Stereological estimates of the subtypes of retrogradely labelled Rb-GFP+ GPe neurons that comprise the pallidostriatal pathway innervating ChAT interneurons. Normalised connectivity for GPe cells expressing FoxP2 or Nkx2-1 with or without co-expression of PV. Asterisks denote **P < 0.01; ns, not significant; one-way ANOVA with Tukey’s post-hoc test. ***K***, Distribution of Rb-GFP+ neurons (dots) across multiple coronal sections of the GPe, shown for each of the cell types examined: FoxP2+ (*left*), Nkx2-1+ PV- (*middle*) and Nkx2-1+ PV+ (*right*). Each dot represents a neuron counted within the 10 µm optical plane of each section. Colours are per animal. The mean centre (‘centre of mass’) was calculated for each of the cell types examined, for each animal (open circles; n=5). Scale bars: ***B***-***C***, 1 mm; ***E***, 20 µm; ***G***, 200 µm; ***H***, 10 µm.

To assess the selectivity of our viral tracing strategy we examined the expression of ChAT in starter neurons in the striatum of injected ChAT-Cre mice (Fig 6E). We found that neurons expressing V5+ and Rb-GFP+ could also express ChAT (Fig 6E). We quantified this co-expression using stereology (Fig 6F, Table S3). The majority of starter neurons expressed ChAT (90.9% ± 3.6%; n=5; Fig 6F), consistent with the molecular phenotype of cholinergic interneurons.

Retrogradely labelled Rb-GFP neurons were localized within the GPe of ChAT-Cre injected animals (Fig 6G). To establish which cell types were present, we tested for some of the known makers of pallidal cell types (Fig 6H-J). Rb-GFP neurons were shown to co-express markers of arkypallidal and prototypic neurons, providing evidence that both these subsets for synaptic connections with cholinergic interneurons (Fig 6H-J).

We estimated the total numbers of striatal starter neurons and Rb-GFP+ GPe neurons in each ChAT-Cre mouse (Fig 6K). The average estimated total number of starter neurons per mouse was 2,052 (± 339; n=5; Table S3) and the average estimated total number of Rb-GFP+ GPe neurons was 2,684 (± 184.5; n=5; Table S4). The average overall connectivity index in the ChAT-Cre mice was 1.40 (± 0.16; n=5) GPe Rb-GFP cells per starter cell.

Analyses of the relative connectivity indexes for each cell type revealed that, for cholinergic interneurons, the most numerous GPe cell type was FoxP2+ neurons (Fig 6L), constituting 55.6% (± 2.6%; n=5) of the estimated neurons. The next most numerous cell type was Nkx2-1 neurons that do not co-express PV (21.1% ± 2.1%; n=5) and then Nkx2-1 neurons co-expressing PV (14.9% ± 1.7%; n=5; Fig 6L). As seen with other cell types, a proportion of neurons did not show any immunoreactivity for any of the markers that we tested (GFP only; 8.4% ± 1.2%; n=5; Fig 6L). Arkypallidal neurons, expressing FoxP2, were found to be significantly more connected with cholinergic interneurons compared to Nkx2-1+ neurons expressing PV (p= 0.0089; Kruskal-Wallis with Dunn’s post hoc test; n=5; Fig 6M).

The coordinates (*x*, *y*) of each retrogradely labelled Rb-GFP neuron in the GPe, within each section, were exported and categorized by cell type for each animal (Fig 6N; n=5; coloured per animal). The ‘centre of mass’ for each cell type was then calculated (Fig 6N). As in the striatum (Fig 6D) the averaged centre for each animal, and for each cell type, was located central within the GPe, and did not show any bias in the medial-lateral or ventral-dorsal planes.

### Selective innervation of striatal cell types by distinct sets of pallidal neurons

Having established the connectivity to specific populations of the major striatal neuronal cell types we can now compare the innervation from distinct subsets of pallidal neurons. If we assess the arkypallidal innervation to SPNs, we find that the normalized connectivity is slightly higher for iSPNs (0.27 ± 0.04) than for dSPNs (0.19 ± 0.01), however this is not significant (Fig 7A; p=0.0857; unpaired t-test). Arkypallidal innervation of striatal interneurons showed that the highest normalized connectivity was with cholinergic interneurons (0.76 ± 0.07), compared to PV interneurons (0.41 ± 0.06; p=0.0077) and SOM interneurons (0.29 ± 0.04; p=0.0022; one-way ANOVA with Tukey’s post hoc test; Fig 7A).

**Figure 7.**
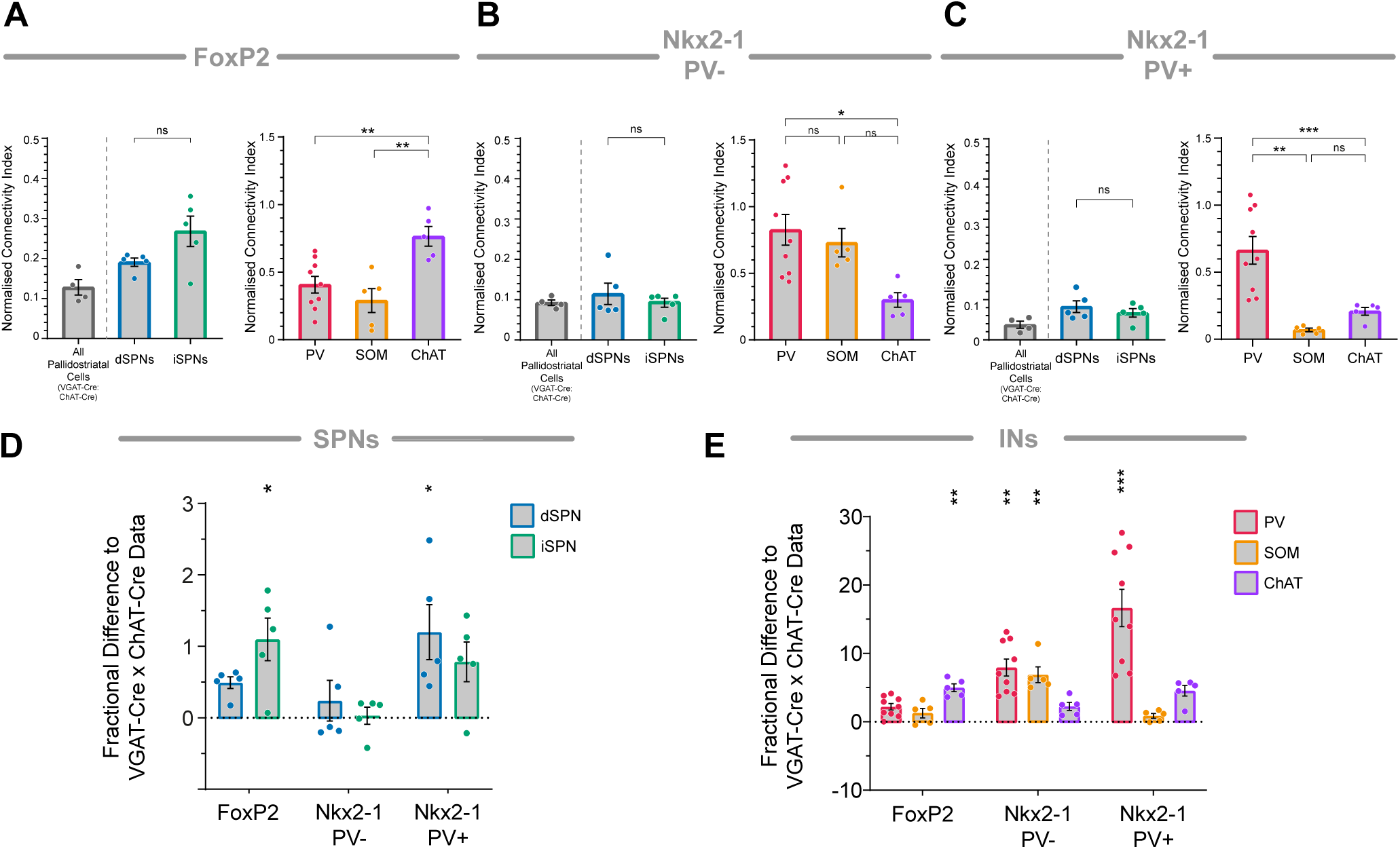
Selective innervation of striatal neurons by subsets of pallidal neurons. ***A***, FoxP2+ Arkypallidal neurons project to direct and indirect pathway SPNs (left) and show a selective preference for innervating ChAT+ cholinergic interneurons (right). ***B***, Nkx2-1+/PV-prototypic neurons innervate both direct and indirect pathway SPNs (left), and to all striatal interneuron classes (right) with a preference for GABAergic interneurons. ***C***, Nkx2-1+/PV+ innervate direct and indirect pathway SPNs, and all classes of striatal interneuron, but with a strong preference for PV interneurons. Quantification of the overall pallidostriatal pathway, to all neuron types in the VGAT-Cre:ChAT-Cre mice, allowed us to calculate a *baseline* projection to which we can compare the relative projections to specific striatal subtypes. ***D***, FoxP2 neurons show an enriched innervation of iSPNs compared to the baseline. NKx2-1-only neurons do not show a selective innervation of a projection neuron type. Nkx2-1/PV+ neurons have a relatively increase innervation of dSPNs. ***E***, For interneurons, FoxP2 arkypallidal neurons have an enriched innervation of ChAT neurons compared to baseline, Nkx2-1+/PV-neurons preferentially innervate GABAergic SOM and PV interneurons and NK2-1+/PV+ neurons show a strong preference for PV interneurons relative to the overall pallidostriatal projection. Asterisks denote ***P < 0.001; **P < 0.01; *P < 0.05; ns, not significant.

Comparison of prototypic innervation to SPNs by neurons expressing Nkx2-1 but not PV, showed that the connectivity is similar between dSPNs (0.11 ± 0.03) and iSPNs (0.095 ± 0.01; ns, p>0.99; Mann-Whitney test). Innervation of interneurons revealed that the greatest connectivity was with PV interneurons (0.83 ± 0.12) and SOM neurons (0.73 ± 0.11) and to a lesser degree with cholinergic interneurons (0.3 ± 0.055; Fig 7B). Indeed, Nkx2-1-only prototypic neurons are more highly connected with PV interneurons, compared to cholinergic interneurons (p=0.0102, one-way ANOVA with Tukey’s post hoc test; Fig 7B).

Prototypic neurons expressing both Nkx2-1 and PV also showed a very similar innervation to dSPNs (0.08 ± 0.01) and iSPNs (0.07 ± 0.01; Fig 7C). Prototypic neurons expressing PV showed the greatest connectivity with striatal PV interneurons (0.66 ± 0.1) when compared with cholinergic neurons (0.21 ± 0.03; p= 0.0040) and SOM interneurons (0.07 ± 0.01; p= 0.0006; one-way ANOVA with Tukey’s post hoc test; Fig 7C).

Since we examined the pallidostriatal pathway, *as a whole*, using the VGAT-Cre:ChAT-Cre mice (Fig 1), we have the overall connectivity data for each GPe cell type. We defined this data as the ‘baseline’ so that we could examine the innervation to each population of striatal cell-type relative to this by calculating the ‘fractional difference’ to baseline. This can give us an indication as to whether particular cell-type specific connections are enriched when compared with the overall projection (Fig7 D,E). To allow for comparisons between cell-types, we accounted for topographical differences by mapping the starter neuron and projection neuron populations (Fig S7). We found that the centre of all starter neuron populations was central within the dorsal striatum for each animal (Fig S7 A,C) and that there does not appear to be any systematic bias in the topography of the pallidostriatal protection, when examined for each cell-type (Fig S7 B,D).

Comparison of GPe connectivity to baseline for SPNs showed that the connection between FoxP2 neurons and iSPNs is significantly enriched (p=0.0109, Kruskal-Wallis with Dunn’s post-hoc test; Fig 7D). There was no significant increase/decrease in the projection from Nkx2-1 prototypic neurons not expressing PV to either dSPNs or iSPNs compared to baseline (Fig 7D). Prototypic neurons expressing Nkx2-1 and PV showed an enriched connectivity with dSPNs (p=0.0473, Kruskal-Wallis with Dunn’s post-hoc test; Fig 7D). Taken together these data suggest a selective projection from arkypallidal neurons to iSPNs and a selective innervation of dSPNs by PV-expressing prototypic neurons (Fig 7D).

Examination of the arkypallidal FoxP2 innervation of striatal interneurons, compared with the baseline pallidostriatal connection, revealed that the connection with cholinergic interneurons was relatively increased (p=0.0014, Kruskal-Wallis with Dunn’s post-hoc test; Fig 7E). This was not the case for either PV or SOM interneurons. Prototypic neurons, not expressing PV, showed the opposite pattern, with a selectively increased innervation of GABAergic PV and SOM interneurons (p= 0.0017 and p=0.009, respectively, Kruskal-Wallis with Dunn’s post-hoc tests; Fig 7E) but not cholinergic interneurons. PV-expressing prototypic neurons show a selectively increased connectivity with striatal PV interneurons (p=0.0004, Kruskal-Wallis with Dunn’s post-hoc test; Fig 7E).

## Discussion

Our results provide a detailed and quantitative map of the pallidostriatal pathway, revealing distinct patterns of connectivity between molecularly defined subtypes of GPe neurons and specific molecular (and functional) subtypes of striatal neurons. Using cell-type-specific monosynaptic rabies tracing in combination with stereology and molecular profiling, we demonstrate that both prototypic and arkypallidal neurons make direct synaptic contacts with all major classes of striatal neurons, including projection neurons and interneurons. However, the selectivity of these connections varies considerably depending on the identity of both the pallidal neurons and the striatal neurons involved.

Consistent with previous studies (Bevan *et al*. 1998; Mallet *et al*. 2012; Abdi *et al*. 2015; Fujiyama *et al*. 2016; Glajch *et al*. 2016; Saunders, Huang and Sabatini 2016; Baker *et al*. 2023; Nambu and Chiken 2024), our findings confirm that arkypallidal neurons are the predominant pallidal cell type innervating the striatum, followed by PV-negative (Nkx2-1+/PV-) prototypic neurons, while the PV-positive (Nkx2-1+/PV+) prototypic neurons constitute <15% of the pathway.

Arkypallidal neurons have been shown to innervate SPNs, but there is limited or inconsistent data on the relative innervation of direct vs. indirect SPNs (Guo *et al*. 2015; Glajch *et al*. 2016; Mallet *et al*. 2016). Here we show that arkypallidal neurons exhibit an enriched innervation of iSPNs. Conversely, PV-prototypic neurons innervate both SPN types without clear selectivity, while PV+ prototypic neurons preferentially target dSPNs, in agreement with previous findings (Saunders, Huang and Sabatini 2016).

Consistent with previous observations (Mallet *et al*. 2012; Klug *et al*. 2018) we demonstrate that pallidal neurons innervate striatal cholinergic interneurons. Our results further reveal that it is arkypallidal neurons that preferentially target cholinergic interneurons and that they are the preferential target of arkypallidal neurons over other striatal interneuron types. In contrast, prototypic pallidal neurons show an enriched innervation of GABAergic interneurons. In addition, the PV-negative prototypic neurons innervate both PV-positive and SOM-positive GABA interneurons to a similar extent, whereas PV-expressing prototypic neurons show a strong preference for striatal PV-positive GABA interneurons, in agreement with prior data (Saunders, Huang and Sabatini 2016).

### Functional Implications

The selective innervation patterns revealed in this study demonstrate that the pallidostriatal pathway is not a uniform projection but rather comprises multiple parallel circuits linking specific subtypes of GPe neurons to specific subtypes of striatal neurons. This organizational principle implies that pallidal neurons, through specialised synaptic connections, can differentially shape striatal microcircuits and, by extension, motor behaviour.

The selective innervation of iSPNs and cholinergic interneurons by arkypallidal neurons raises an apparent paradox: if arkypallidal neurons inhibit iSPNs, one might predict facilitation of movement, since activity in the indirect pathway is typically associated with a decrease in motor output - however arkypallidal neuron activation is associated with suppression of actions (Mallet *et al*. 2016; Pamukcu *et al*. 2020; Ketzef and Silberberg 2021; Baker *et al*. 2023; Giossi *et al*. 2024). One possible explanation lies in the enriched connectivity to cholinergic interneurons. These interneurons powerfully influence striatal dopamine signalling and SPN excitability, strongly modulating the networks contributing to motor function, learning, reinforcement, and behavioural flexibility (Zhou, Wilson and Dani 2002). Through this pathway, arkypallidal neurons may shift the dopamine-acetylcholine balance in the striatum, indirectly altering SPN output and modulating the likelihood of movement initiation or suppression. Moreover, arkypallidal neurons receive prominent input from dSPNs (Cui *et al*. 2021a; Ketzef and Silberberg 2021), and the cortex (Karube *et al*. 2019; Abecassis *et al*. 2020) positioning them to integrate goal-directed signals with feedback from the direct pathway. Additional inputs from the STN and iSPNs can strongly influence arkypallidal neuron activity through a di-synaptic mechanism involving prototypic neuronal axon collaterals (Aristieta *et al*. 2021). These convergent inputs position arkypallidal neurons as integrative hubs, capable of selectively inhibiting spatially and temporally defined subsets of striatal neurons to cancel unwanted or inappropriate movements.

In contrast, prototypic GPe neurons, particularly those expressing PV, show selective innervation of striatal GABAergic interneurons, especially PV+ striatal interneurons as well as dSPNs. This aligns with prior data indicating that PV+ prototypic neurons exert strong, fast GABAergic inhibition of fast-spiking and NPY interneurons, but with only a minimal impact on SPNs (Saunders, Huang and Sabatini 2016). The preferential targeting of inhibitory interneurons suggests a disinhibitory mechanism for modulating striatal projection neuron excitability, potentially contributing to rapid gating or synchronization of activity during motor execution.

Parvalbumin-negative prototypic neurons selectively target both PV+ and SOM+ interneurons and show comparatively less selectivity for SPNs. Their broader interneuron engagement suggests they may exert more diffuse inhibitory/disinhibitory control, tuning striatal responsiveness across multiple interneuron-defined microcircuits. This connectivity may be important for synchronisation of striatal activity across motor tasks.

Taken together, the selective organization of the pallidostriatal pathway points to a functional division of labour: arkypallidal neurons modulate striatal output through iSPNs and cholinergic interneurons, influencing motivational and contextual aspects of behaviour, whereas prototypic neurons modulate the temporal precision and excitability of local networks via GABAergic interneurons. Understanding how these pathways interact during behaviour will be critical for interpreting the contributions of GPe subtypes to movement control and dysfunction in disease.

### Technical Limitations

Although the present study provides direct evidence of synaptic connectivity between populations of pallidal and striatal neurons, there are, of course, limitations. First, although rabies tracing allows for selective and robust anatomical mapping of monosynaptic connections, it does not offer insight into the synaptic efficacy nor subcellular localisation of the connections identified. We cannot determine the number of synapses formed by each presynaptic neuron, the precise location of those synapses on the postsynaptic cell, or the extent of convergence and divergence across the network. Second, although we characterised the major known GPe cell types using established molecular markers, the cellular taxonomy of the GPe remains an area of active investigation. There is currently no field-wide consensus on the full extent of pallidal neuron diversity; and overlap between molecular markers introduces additional ambiguity. Our work therefore cannot capture all reported subtypes, findings leave open the possibility that additional GPe cell types contribute to the pallidostriatal projection, or that further specificity exists within the populations profiled here. Third, even with rigorous stereological quantification and topographical control, local circuit interactions within both the GPe and striatum likely modulate how these projections influence downstream computation. Thus, anatomical specificity must be interpreted within the context of dynamic circuit processing.

### Future Directions

This study establishes a foundational anatomical framework for understanding how the GPe shapes striatal function through distinct circuit motifs. Future work combining circuit-specific functional manipulations with behavioural paradigms and *in vivo* recordings will be essential to dissect how these selective pathways contribute to action selection, suppression, and learning. Furthermore, understanding how these pallidostriatal networks are altered in disease states, such as Parkinson’s or compulsive disorders, could uncover new therapeutic targets within the basal ganglia circuitry.

## Methods

### Animals

Experimental procedures were performed on mice and were conducted at the University of Oxford in accordance with the Animals (Scientific Procedures) Act, 1986 (United Kingdom).

Male and female mice aged from 2.5 to 8 months old were used for all experiments. Seven lines of genetically altered mice were used; all were bred to a C57Bl6/J background, and only mice heterozygous/hemizygous for the transgene(s) were used in experiments. To retrogradely label GPe neurons innervating striatal neurons as a whole, we used a ‘double transgenic’ line produced by crossing **VGAT-Cre** mice (*Slc32a1^tm2(cre)Lowl^*/J; Jackson Laboratory; RRID:IMSR_JAX:016962) with **ChAT-Cre mice** (B6;129S6-*Chat^tm2(cre)Lowl^*/J; Jackson Laboratory; RRID:IMSR_JAX:006410). To retrogradely label pallidal neurons innervating more restricted populations of neurons in striatum, we used the following lines: **Drd1a-Cre** mice (B6.FVB(Cg)-Tg(Drd1-cre)EY262Gsat/Mmucd; GENSAT/MMRRC; RRID:MMRRC_030989-UCD); **Adora2a-Cre** mice (B6.FVB(Cg)-Tg(Adora2a-cre)KG139Gsat/Mmucd; GENSAT/MMRRC; RRID:MMRRC_036158-UCD); **PV-Cre** mice (B6;129P2-Pvalbtm1(cre)Arbr/J; Jackson Laboratory; RRID:IMSR_JAX:008069); **SOM-Cre mice** (Ssttm2.1(cre)Zjh/J; Jackson Laboratory; RRID:IMSR_JAX:013044); and **ChAT-Cre** mice (B6;129S6-Chattm2(cre)Lowl/J; Jackson Laboratory; RRID:IMSR_JAX:006410).

### Viruses

To carry out monosynaptic retrograde tracing from neurons in the dorsal striatum, we made sequential use of a single Cre-dependent ‘helper virus’ (AAV5-DIO-TVA^V5^-RG; bicistronically expressing TVA receptor fused to a V5 tag, and the rabies glycoprotein [RG]) and a ‘modified rabies virus’ ([EnvA]-SADΔG-EGFP; pseudotyped with EnvA, RG deleted, and expressing enhanced GFP (Ährlund-Richter *et al*. 2019; Kondabolu *et al*. 2023).

### Stereotaxic intracerebral injections of rhabdovirus vectors

General anaesthesia was induced and maintained with isoflurane (1.0-3.0% v/v in O_2_). Animals received perioperative analgesic (buprenorphine, 0.1 mg/kg, s.c.; Ceva) and were placed in a stereotaxic frame (Kopf Instruments). Incision margins were first infiltrated with local anaesthetic (0.5% w/v bupivacaine [Marcaine]; Aspen). Body temperature was maintained at ∼37°C by a homeothermic heating device (Harvard Apparatus). In the first step, we used a glass micropipette to unilaterally inject 60-120 nl of helper virus into the central aspects of the dorsal striatum of Cre-expressing mice, using the following coordinates (caudal approach at an angle of 20° to vertical): 0.00 mm anterior of Bregma, 2.20 mm lateral of Bregma, and 2.70 mm ventral to the brain surface. To minimize reflux, the micropipette was left in place for ∼10 min after the injection. Allowing 21 d for neuron transduction, Cre-mediated recombination, and the generation of ‘starter’ neurons expressing all components necessary for retrograde labelling of their presynaptic partners, we then as a second step injected 60-120 nl of modified rabies virus into the same striatal locations, using the following coordinates (vertical, no angle): 1.00 mm anterior of Bregma,

2.20 mm lateral of Bregma, and 2.50 mm ventral to the brain surface. To minimize reflux, the micropipette was left in place for ∼10 min after the injection. Mice were maintained for 7 d after surgery to allow for starter neuron infection and retrograde trans-synaptic labelling of neurons providing monosynaptic inputs to starters. Thereafter, mice were killed with pentobarbitone (1.5 g/kg, i.p.; Animalcare) and transcardially perfused with 20-50 ml of 1×PBS, pH 7.4 (PBS), followed by 30-100 ml 4% w/v PFA in 0.1 M phosphate buffer, pH 7.4 (PB). Brains were removed and left overnight in fixative at 4°C before sectioning.

### Tissue processing and indirect immunofluorescence

Brains were embedded in agar (3-4% w/v dissolved in dH_2_O) before being cut into 50 µm coronal sections on a vibrating microtome (VT1000S; Leica Microsystems). Free-floating tissue sections were collected in series, washed in PBS, and stored in PBS containing 0.05% w/v sodium azide (Sigma) at 4°C until processing for indirect immunofluorescence to reveal molecular markers (Abdi et al., 2015). Briefly, after 1 h of incubation in “Triton PBS” (PBS with 0.3% v/v Triton X-100 and 0.02% w/v sodium azide) containing 10% v/v normal donkey serum (NDS; 017-000-121, Jackson ImmunoResearch Laboratories, RRID:AB_2337258), sections were incubated overnight at room temperature, or for 72 h at 4°C, in Triton PBS containing 1% v/v NDS and a mixture of between two and four primary antibodies (Table S1). After exposure to primary antibodies, sections were washed in PBS and incubated overnight at room temperature in Triton PBS containing an appropriate mixture of secondary antibodies (all raised in donkey) with minimal cross-reactivity and that were conjugated to the following fluorophores: AMCA (1:250 dilution; Jackson ImmunoResearch Laboratories); Alexa Fluor 488 (1:1000; Invitrogen); Cy3 (1:1000; Jackson ImmunoResearch Laboratories), Cy5 or DyLight 647 (1:500; Jackson ImmunoResearch Laboratories).

To optimize immunolabeling for some of the primary antibodies we used heat pre-treatment as a means of antigen retrieval (Table S1) (Mallet *et al*. 2012; Garas *et al*. 2016, 2018; Sharott *et al*. 2017). Following incubation of sections in primary antibodies (that did not require antigen retrieval) then secondary antibodies, sections were sequentially washed in PBS and citrate buffer (10 mM citric acid, pH=6) before incubation in citrate buffer at 80°C for 1 h. Sections were then allowed to come to room temperature, and washed back into PBS, before incubation overnight at room temperature in PBS containing 1% v/v NDS and primary antibodies (Table S1). Sections were then washed in PBS and incubated overnight at room temperature in PBS containing an appropriate mixture of secondary antibodies (as above).

After binding of primary and secondary antibodies, and final washing in PBS, sections were mounted on glass slides and cover-slipped using Vectashield Mounting Medium (H-1000, Vector Laboratories, RRID:AB_2336789) or SlowFade Diamond Antifade Mountant (S36972, ThermoFisher Scientific). Coverslips were sealed using nail varnish and slides stored at 4°C before imaging.

### Stereological quantification for retrograde labelling experiments in vivo

A version of design-based stereology, the ‘modified optical fractionator’, was used to generate unbiased cell counts and determine the proportions of a given population of neurons that expressed certain combinations of molecular markers (Abdi *et al*. 2015; Dodson *et al*. 2015; Garas *et al*. 2016, 2018; Kondabolu *et al*. 2023). All stereology, imaging for stereology, and cell counting, was performed using Stereo Investigator software (v. 2019.1.4, MBF Bioscience, RRID:SCR_002526). Acquisition of images for stereological sampling was carried out on an AxioImager.M2 microscope (Zeiss) equipped with an ORCA Flash-4.0 LT digital CMOS camera (Hamamatsu), an Apotome.2 (Zeiss), and a Colibri 7 LED light source (Type R[G/Y]B-UV, Zeiss). Appropriate sets of filter cubes were used to image the fluorescence channels: AMCA (excitation 299–392 nm, beamsplitter 395 nm, emission 420-470 nm); Alexa Fluor 488 (excitation 450–490 nm, beamsplitter 495 nm, emission 500-550 nm); Cy3 (excitation 532-558 nm, beamsplitter 570 nm, emission 570-640 nm); and Cy5/DyLight 649 (excitation 625-655 nm, beamsplitter 660 nm, emission 665-715 nm). Images of each of the channels were taken sequentially and separately to negate possible crosstalk of signal across channels. To quantify retrogradely-labelled GPe neurons, we first defined (using a 10× objective lens; 0.45 NA; Plan-Apochromat, Zeiss) the borders of the GPe according to the expression of PV (Abdi *et al*. 2015); the full extent of the GPe was imaged for each series examined in each mouse. To quantify striatal starter neurons, we first defined (using a 10× objective) the outer boundaries of regions within striatum that contained neurons immunoreactive for V5, an indicator of neurons transduced with the helper virus, and thus, potential starter cells (Kondabolu *et al*. 2023). After delineating the borders of GPe and the boundaries of striatal regions containing starter neurons, images for stereological sampling were acquired using the optical fractionator workflow in Stereo Investigator, employing a 2 µm-thick ‘guard zone’, and an unbiased 2D counting frame and grid frame of 600 × 600 µm (i.e., 100% of the region in the X, Y plane was sampled). Z-stacked images across a 10 µm-thick ‘optical disector’ were acquired using a 20× objective lens (0.8 NA; Plan-Apochromat), and images or ‘optical sections’ were taken in 1 µm steps to ensure no loss of signal in the Z-axis. The captured images were used for counting offline. A neuron was only counted when its nucleus came into sharp focus within the disector; neurons with nuclei already in focus in the top optical section of the disector were ignored. The use of stereology, and this optical disector probe in particular, ensured that we could generate robust and unbiased cell counts in a timely manner. For a given molecular marker, X, we designate positive immunoreactivity (confirmed expression) as X+, and undetectable immunoreactivity (no expression) as X-. A neuron was classified as not expressing the tested molecular marker only when positive immunoreactivity could be observed in other cells on the same optical section as the tested neuron. Striatal starter neurons were defined by their co-expression of immunoreactivity for V5 and rabies-encoded GFP (Rb-GFP). To ensure a high level of precision in the cell counts, data was only included from individual mice when the Coefficient of Error (CE; using the Gundersen method) was ≤ 0.1 with a smoothness factor of m = 1 (West, Slomianka and Gundersen 1991; Gundersen *et al*. 1999; West 1999, 2012). The CE provides an estimate of sampling precision, which is independent of biological variance. As the value approaches zero, the uncertainty in the estimate precision reduces. The number of sections counted per mouse was thus dependent on variability; sections/series were added to the analysis until the CE ≤ 0.1 for each animal. Images for figures were acquired with a confocal microscope (LSM880, Zeiss). All image adjustments were applied to every pixel.

### Topographical analysis of virally labelled neurons

To evaluate and compare the distributions of virally labelled neurons across animals and mouse lines, the relative two-dimensional coordinates of individual virus-labelled neurons in the striatum and GPe, as well as the contours of both the structures, were exported in the XML format from Stereo Investigator (v. 2019.1.4, MBF Bioscience, RRID:SCR_002526) and processed with MATLAB software (MathWorks, version: R2021b) using custom functions and scripts (https://github.com/kouichi-c-nakamura/doig-gpe-striatum-trace-mapping) in the same way as previously described (Garas *et al*. 2016). For each coronal section, the *x* and *y* (mediolateral and dorsoventral) coordinates of the virally labelled neurons were normalised to the encompassing contours of the striatum or GPe so that they range from −1 to +1; The coordinate (0, 0) for a striatal or GPe cell means that the cell was positioned equidistantly from the mediodorsal and dorsoventral limits of the striatum or GPe. Although this method does not consider the changes in the size and shape of striatum or GPe along the anteroposterior axis, it achieved fair alignment of coronal sections as indicated by the overlaid contours (see Supplementary Fig. 7). For the graphical representations of neuronal positions, the average aspect ratios of the striatum and GPe were computed and used to draw the structures. The centre of mass of a group of neurons was computed as a numeric mean of the normalized coordinate values of the constituent neurons and plotted to represent the group.

### Statistics and Reproducibility

For experiments entailing monosynaptic retrograde tracing from neurons in the striatum, our use of differently angled stereotaxic injections of helper virus and modified rabies virus helped ensure that starter neurons were restricted to dorsal striatum. As a control for the Cre- and TVA-dependence of neuronal labelling, we injected helper virus and/or modified rabies virus into the striata of adult wild-type mice (*n* = 2). As previously reported for the vectors we use here (Ährlund-Richter *et al*. 2019; Kondabolu *et al*. 2023), these injections resulted in negligible numbers of starter neurons and retrogradely labelled neurons.

For each experiment, descriptions of critical variables (e.g., number of mice, neurons, and other samples evaluated) as well as details of statistical design and testing can be found in the Results and Supplementary Tables S2-S4. For monosynaptic retrograde tracing experiments, the ratio (*x*/*y*) of the stereologically-estimated total number of striatal starter neurons (*x*) to the estimated total number of retrogradely-labelled GPe neurons (*y*) was calculated for each animal (Do et al., 2016; Choi et al., 2019) and is referred to as the ‘normalized connectivity index’. In order to compare the connectivity data for specific subpopulations of striatal neurons relative to the overall pallidostriatal projection we defined the data from VGAT-Cre:ChAT-Cre mice as the ‘baseline.’ The ‘fractional difference’ was then calculated as [(value – baseline)/baseline]. Graphing and statistical analyses were performed with GraphPad Prism (v9, RRID:SCR_002798). If datasets were normally distributed with equal variance, then parametric statistical tests were used, and non-parametric tests used in all other cases; all statistical tests are indicated in the text. The Shapiro-Wilk test was used to judge whether datasets were normally distributed (*p* ≤ 0.05 to reject) and the Brown-Forsythe test was used to test for the equality of group variances. Significance for all statistical tests was set at *p* < 0.05 (exact *p* values are given in the text). Data are represented as group means ± SEMs unless stated otherwise, with some plots additionally showing individual samples (mice or neurons) as appropriate.

## Data Availability

The authors declare that all relevant data are included in the paper and its supplementary information files.

## Code Availability

All custom code used in this study is freely available here: https://github.com/kouichi-c-nakamura/doig-gpe-striatum-trace-mapping

## Acknowledgements

The work of N.M.D., C.M., K.N., and P.J.M. was supported by the UK Medical Research Council (Awards MC_UU_12024/2 and MC_UU_00003/5 to P.J.M. and MC_UU_00003/6 to AS and the Wellcome Trust (Investigator Award 101821 to P.J.M.). The work of K.M. was supported by the Swedish Research Council and the Swedish Brain Foundation (Hjärnfonden). We thank J. Westcott, L. Conyers, H. Zhang, and B. Micklem, for excellent technical assistance. We thank J.P. Bolam for comments on early versions of the manuscript.

## Author Contributions

N.M.D and P.J.M conceived and planned the experiments. N.M.D and C.M carried out the experiments. N.M.D, K.N and A.S analysed the data. N.M.D, K.N, A.S, K.M & P.J.M contributed to the interpretation of the results. N.M.D and P.J.M wrote the manuscript. All authors provided critical feedback and helped shape the research, analysis, and manuscript.

