## Supplementary material for "Selective innervation of subpopulations of striatal neurons by distinct sets of neurons of the external globus pallidus": Tables_S1-S4_Figure_S7

### Supplementary Information

#### *Table S1. Antibody Details*

| Antigen | Species | Company | Catalogue Number | Dilution | Antibody ID (RRID) | Heat Induced Antigen Retrieval | References |
|---|---|---|---|---|---|---|---|
| **Choline Acetyl Transferase (ChAT)** | Goat | Millipore | AB144P | 1 in 500 | AB_2079751 | N | Abdi et al., 2016; Garas et al., 2016; Kondabolu et al., 2023; Sharott et al., 2012, 2017. |
| **COUP TF1-interacting protein 2 (Ctip2)** | Rat | Abcam | ab18465 | 1 in 500 | AB_2064130 | Y | Garas et al., 2018; Garas et al., 2016 |
| **Forkhead Box protein 2 (FoxP2)** | Goat | Santa Cruz | sc-21069 | 1 in 500 | AB_2107124 | Y | Kondabolu et al., 2023; Abdi et al., 2016; Dodson et al., 2015 |
| **Green Fluorescent Protein (GFP)** | Mouse | Invitrogen | A-11120 | 1 in 500 | AB_221568 | N | Kesner et al., 2021; Chung et al., 2017 |
| **Green Fluorescent Protein (GFP)** | Rat | Nacalai Teaque | 04404-84 | 1 in 1000 | AB_10013361 | N | Kondabolu et al., 2023; Dodson et al., 2015 |
| **Nitric Oxide Synthase (NOS)** | Goat | Abcam | ab1376 | 1 in 500 | AB_300614 | N | Kondabolu et al., 2023; Matsuda et al., 2020 |
| **NK2 Homeobox 1 (Nkx2-1)** | Rabbit | Santa Cruz | sc-13040 | 1 in 500 | AB_793532 | N | Dodson et al., 2015 |
| **Parvalbumin (PV)** | Guinea-Pig | Synaptic Systems | 195004 | 1 in 1000 | AB_2156476 | N | Garas et al., 2016; Sharott et al., 2012 |
| **Preproenkephalin (PPE)** | Rabbit | LifeSpan BioSciences | LS-C23084 | 1 in 5000 | AB_902714 | Y | Sharott et al., 2017; Garas et al., 2016; Dodson et al., 2015 |
| **Somatostatin (SOM)** | Mouse | GeneTex | GTX71935 | 1 in 250 | AB_383280 | N | Zhu et al., 2020; Yuan et al., 2017 |
| **V5** | Chicken | Abcam | ab9113 | 1 in 2000 | AB_307022 | N | Kondabolu et al., 2023; Lazaridis et al., 2019 |

***Table S2. Stereological Parameters***

| | Region | n | Total Number of Neurons Counted | | Number of Sections | | Number of Sampling Sites | | Coefficient of Error (Gundersen; m=1) | |
|---|---|---|---|---|---|---|---|---|---|---|
| | | | **Mean** | **SEM** | **Mean** | **SEM** | **Mean** | **SEM** | **Mean** | **SEM** |
| **VGAT-Cre:ChAT-Cre** | Striatum | 4 | 1263.75 | 141.00 | 6.50 | 0.29 | 101.75 | 9.71 | 0.03 | 0.002 |
| **Drd1a-Cre** | Striatum | 5 | 510.20 | 133.69 | 6.00 | 0.45 | 46.60 | 5.80 | 0.04 | 0.007 |
| **Adora2a-Cre** | Striatum | 5 | 386.00 | 104.40 | 5.80 | 0.20 | 40.60 | 4.53 | 0.05 | 0.006 |
| **PV-Cre** | Striatum | 9 | 39.70 | 9.10 | 8.20 | 1.1 | 67.89 | 5.60 | 0.08 | 0.004 |
| **SOM-Cre** | Striatum | 5 | 103.60 | 18.30 | 7.40 | 1.7 | 58.60 | 12.90 | 0.06 | 0.008 |
| **ChAT-Cre** | Striatum | 5 | 102.60 | 16.90 | 7.00 | 0.40 | 78.00 | 3.21 | 0.05 | 0.013 |
| **VGAT-Cre:ChAT-Cre** | GPe | 4 | 345.50 | 47.61 | 7.00 | 0.00 | 46.75 | 1.11 | 0.06 | 0.005 |
| **Drd1a-Cre** | GPe | 5 | 206.80 | 34.40 | 7.80 | 0.20 | 44.40 | 0.87 | 0.07 | 0.005 |
| **Adora2a-Cre** | GPe | 5 | 169.20 | 33.50 | 7.20 | 0.40 | 42.80 | 2.52 | 0.08 | 0.007 |
| **PV-Cre** | GPe | 9 | 117.00 | 14.20 | 13.90 | 0.4 | 76.11 | 2.22 | 0.07 | 0.007 |
| **SOM-Cre** | GPe | 5 | 137.6 | 13.5 | 9.8 | 1.8 | 57.4 | 10.5 | 0.09 | 0.004 |
| **ChAT-Cre** | GPe | 5 | 134.40 | 9.20 | 7.00 | 0.30 | 62.40 | 7.19 | 0.09 | 0.004 |

***Table S3. Specificity of Starter Neurons***

| Transgenic Animal | n | Estimated Total Number of Starter Neurons (V5+ GFP+) for each antigen tested | | | | | | | | | |
|---|---|---|---|---|---|---|---|---|---|---|---|
| **VGAT-Cre:ChAT-Cre** | 4 | **Ctip2- ChAT-** | | **Ctip2+ ChAT-** | | **Ctip2- ChAT+** | | | | **Total Starter Neurons** | |
| | | **Mean** | **SEM** | **Mean** | **SEM** | **Mean** | **SEM** | | | **Mean** | **SEM** |
| | | 2795.0 | 530.0 | 21895.0 | 2289.0 | 585.0 | 99.0 | | | 25275.0 | 2820.0 |
| **Drd1a-Cre** | 4 | **Ctip2- PPE-** | | **Ctip2+ PPE-** | | **Ctip2+ PPE+** | | **Ctip2- PPE+** | | **Total Starter Neurons** | |
| | | **Mean** | **SEM** | **Mean** | **SEM** | **Mean** | **SEM** | **Mean** | **SEM** | **Mean** | **SEM** |
| | | 1288.0 | 330.0 | 10275.0 | 3242.0 | 75.0 | 38.6 | 0.0 | 0.0 | 11638.0 | 2996.0 |
| **Adora2a-Cre** | 5 | **Ctip2- PPE-** | | **Ctip2+ PPE-** | | **Ctip2+ PPE+** | | **Ctip2- PPE+** | | **Total Starter Neurons** | |
| | | **Mean** | **SEM** | **Mean** | **SEM** | **Mean** | **SEM** | **Mean** | **SEM** | **Mean** | **SEM** |
| | | 155.0 | 23.7 | 171.0 | 37.3 | 7682.0 | 1988.0 | 255.0 | 159.0 | 8263.0 | 2139.0 |
| **PV-Cre** | 9 | **PV-** | | **PV+** | | | | | | **Total Starter Neurons** | |
| | | **Mean** | **SEM** | **Mean** | **SEM** | | | | | **Mean** | **SEM** |
| | | 34.4 | 11.4 | 605.6 | 98.8 | | | | | 640.0 | 106.9 |
| **SOM-Cre** | 5 | **SOM- NOS-** | | **SOM+ NOS-** | | **SOM+ NOS+** | | **SOM- NOS+** | | **Total Starter Neurons** | |
| | | **Mean** | **SEM** | **Mean** | **SEM** | **Mean** | **SEM** | **Mean** | **SEM** | **Mean** | **SEM** |
| | | 0.0 | 0.0 | 52.0 | 23.3 | 1730.8 | 160.3 | 42.6 | 23.2 | 1825.3 | 142.3 |
| **ChAT-Cre** | 5 | **ChAT-** | | **ChAT+** | | | | | | **Total Starter Neurons** | |
| | | **Mean** | **SEM** | **Mean** | **SEM** | | | | | **Mean** | **SEM** |
| | | 164.0 | 65.8 | 1888.0 | 368.2 | | | | | 2052.0 | 339.0 |

##### *Table S4. Retrogradely Labelled Rb-GFP Neurons in the GPe*

| | Estimated Total Number of GPe Neurons (Rb-GFP+) for each marker tested | | | | | | | | | | |
|---|---|---|---|---|---|---|---|---|---|---|---|
| **Transgenic Animal** | **n** | **GFP+ only** | | **FoxP2+** | | **Nkx2-1+ PV-** | | **Nkx2-1+ PV+** | | **Total** | |
| | | **Mean** | **SEM** | **Mean** | **SEM** | **Mean** | **SEM** | **Mean** | **SEM** | **Mean** | **SEM** |
| **VGAT-Cre:ChAT-Cre** | 4 | 625.7 | 52.6 | 3175.7 | 427.6 | 2312.1 | 223.0 | 941.4 | 201.6 | 7055.0 | 807.7 |
| **Drd1a-Cre** | 5 | 412.0 | 130.0 | 1904.0 | 354.0 | 1048.0 | 168.0 | 772.0 | 124.0 | 4136.0 | 687.0 |
| **Adora2a-Cre** | 5 | 208.0 | 74.2 | 1976.0 | 381.0 | 728.0 | 158.0 | 472.0 | 74.2 | 3384.0 | 669.0 |
| **PV-Cre** | 9 | 122.3 | 21.4 | 230.7 | 37.4 | 457.6 | 57.7 | 366.9 | 62.9 | 1177.6 | 145.9 |
| **SOM-Cre** | 5 | 268.7 | 56.1 | 530.7 | 156.2 | 1308.7 | 172.2 | 127.7 | 26.0 | 2235.7 | 369.9 |
| **ChAT-Cre** | 5 | 228.0 | 41.3 | 1500.0 | 149.8 | 568.0 | 63.7 | 388.0 | 21.5 | 2684.0 | 184.5 |

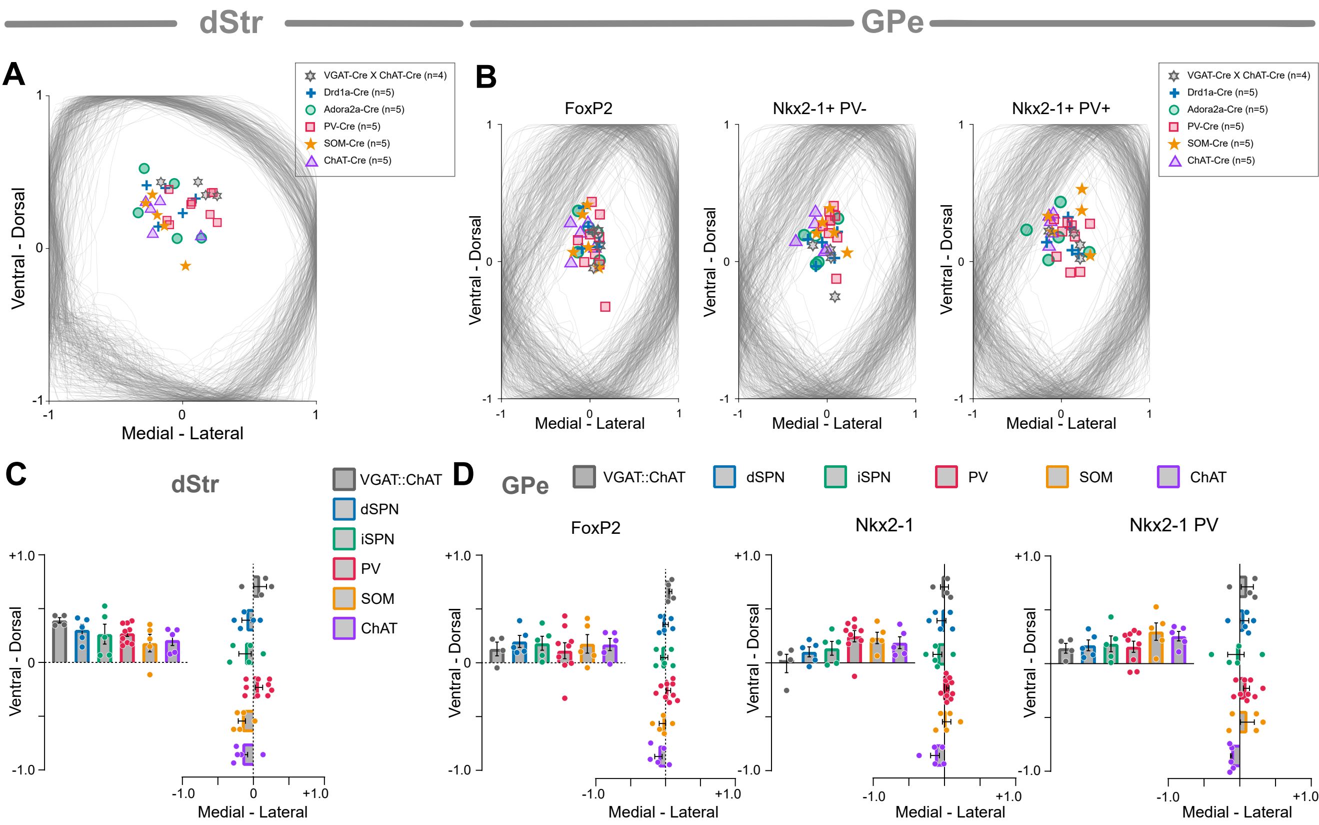

**Supplementary Figure 7. Topographical analyses of striatal and pallidal neurons**
***A***, Overlay of all striatal contours, withing a normalised framework. Each symbol indicates the average centre of mass for striatal starter neurons for each striatal cell type (mouse strain). ***B***, Overlay of pallidal contours for each subset of pallidal neuron, symbols indicate the average centre of mass for counted pallidal neurons within each mouse strain. ***C***, Quantification of striatal starter neuron topography for each cell type in the ventral-dorsal and medial-lateral planes. Data shows that there was no significant bias in the location of the starter neurons. ***D***, Quantification of topographical mapping of pallidal neuron types in ventral-dorsal and medial-lateral planes. Data indicate that there was no significant bias in the pallidal neurons projecting to the striatum. Taken together, these data show that selectivity in pallidostriatal projections are not due to the location of injections or differences in the topography.
